# *Mycobacterium tuberculosis* transcriptional regulator Rv1019 is upregulated in hypoxia and negatively regulates *Rv3230c*-*Rv3229c* operon encoding enzymes in the oleic acid biosynthetic pathway

**DOI:** 10.1101/2022.04.26.489556

**Authors:** Akhil Raj Pushparajan, Lekshmi K Edison, Ramakrishnan Ajay Kumar

## Abstract

The main obstacle in eradicating tuberculosis is the ability of *Mycobacterium tuberculosis* to remain dormant in the host, and then to get reactivated even years later under immuno-compromised conditions. Transcriptional regulation in intracellular pathogens plays an important role in adapting to the challenging environment inside the host cells. Previously, we demonstrated that Rv1019, a putative transcriptional regulator of *M. tuberculosis* H37Rv, is an autorepressor. We showed that, *Rv1019* is cotranscribed with *Rv1020* (*mfd*) and *Rv1021* (*mazG*) encoding DNA repair proteins and negatively regulates the expression of these genes. In the present study, we show that Rv1019 also regulates the expression of the genes *Rv3230c* and *Rv3229c* (*desA3*) which form a two-gene operon in *M. tuberculosis*. Constitutive expression of *Rv1019* in *M. tuberculosis* significantly downregulated the expression of these genes. Employing Wayne’s hypoxia-induced dormancy model of *M. tuberculosis*, we show that *Rv1019* is upregulated (3-fold) under hypoxia. Finally, by reporter assay, using *M. smegmatis* as a model, we validate that Rv1019 is recruited to the promoter of *Rv3230c-Rv3229c* during hypoxia and negatively regulates this operon which is involved in the biosynthesis of oleic acid.

## Introduction

Tuberculosis (TB), caused by *Mtb*, is one of the oldest diseases and is the leading cause of death of humans by an infectious agent [1]. It is estimated by the World Health Organization (WHO) that one-third of the world’s population is infected with *Mtb* causing approximately 1.2 million deaths annually [2].

*Mtb* has the ability to adapt to and persist in the human body in a metabolically inactive state in structures called ‘granuloma’, without manifesting any clinical symptoms [3]. The ability of *Mtb* to remain dormant is one of the major reasons for the ineffectiveness of TB eradication programs, causing longer duration of treatment and emergence of drug resistance [4]. Changes in the physiology of *Mtb* during its transition from actively growing state to dormancy have been studied using genomic, transcriptomic, proteomic and metabolomic approaches [5] [6] [7] [8]. Availability of the complete sequence of *Mtb* genome has led to a better understanding of the pathogen for devising therapeutic strategies [5] [9].

Previously, to study the physiological changes that occur in *Mtb* during dormancy and reactivation, we profiled the proteome of dormant and reactivated bacteria by developing an *in vitro* dormancy-reactivation system employing Wayne’s hypoxia-induced dormancy model [8]. Wayne’s dormancy model has proven to be a very effective and simple method to study the molecular mechanisms in dormant bacteria, and to discover novel therapeutic agents [10]. Also, Wayne’s model is proven to be clinically correlated to human anaerobic latent lesions containing dormant bacilli [11]. Our interest was to dissect the role of transcriptional regulatory proteins which regulate the expression of the genes the products of which help the pathogen adapt to unfavourable hostile conditions. There are 214 transcriptional regulatory proteins annotated to be present in *Mtb* which regulate expression of genes at the level of transcription [12]. Characterization of each of them is critical to unravel the downstream biochemical processes controlled by them.

In our previous study, we characterized Rv1019 [13], a putative transcriptional regulator of *M. tuberculosis* H37Rv which we found to be upregulated in hypoxia-induced dormancy models [8] [14]. We found that under normal aerobic conditions of growth this regulator acts as an autorepressor and forms dimers. We also showed that *Rv1019* is cotranscribed with downstream genes *Rv1020* and *Rv1021* which are involved in DNA repair mechanisms in *Mtb*. In the present study, we made an effort to understand the role of Rv1019 in regulating the expression of two other genes viz. *Rv3230c* and *Rv3229c* (*desA3*) in *Mtb*. Our results show that *Rv3230c* and *Rv3229c* are cotranscribed as an operon and Rv1019 negatively regulates this operon under hypoxia.

## Results

### *Rv3230c* and *Rv3229c* form a two-gene operon in *M. tuberculosis*, and Rv1019 acts as its repressor

Overexpression studies of transcriptional regulators in *Mtb* have shown that Rv1019 could be recruited to the promoter of *Rv3230c* and *Rv3229c* [15] [12]. To verify the *in vivo* binding of Rv1019 on *Rv3230c* and *Rv3229c* promoter, we performed a chromatin immunoprecipitation on the lysate of log-phase *M. tuberculosis* with Rv1019 antibodies and checked the precipitated DNA for *Rv3230c* and *Rv3229c* promoter by PCR. We were able to amplify the promoter of *Rv3230c* but not that of *Rv3229c*, indicating that under normal aerobic laboratory growth conditions Rv1019 can be recruited to *Rv3230c* promoter (**Fig. 1A**).

**Fig. 1.**
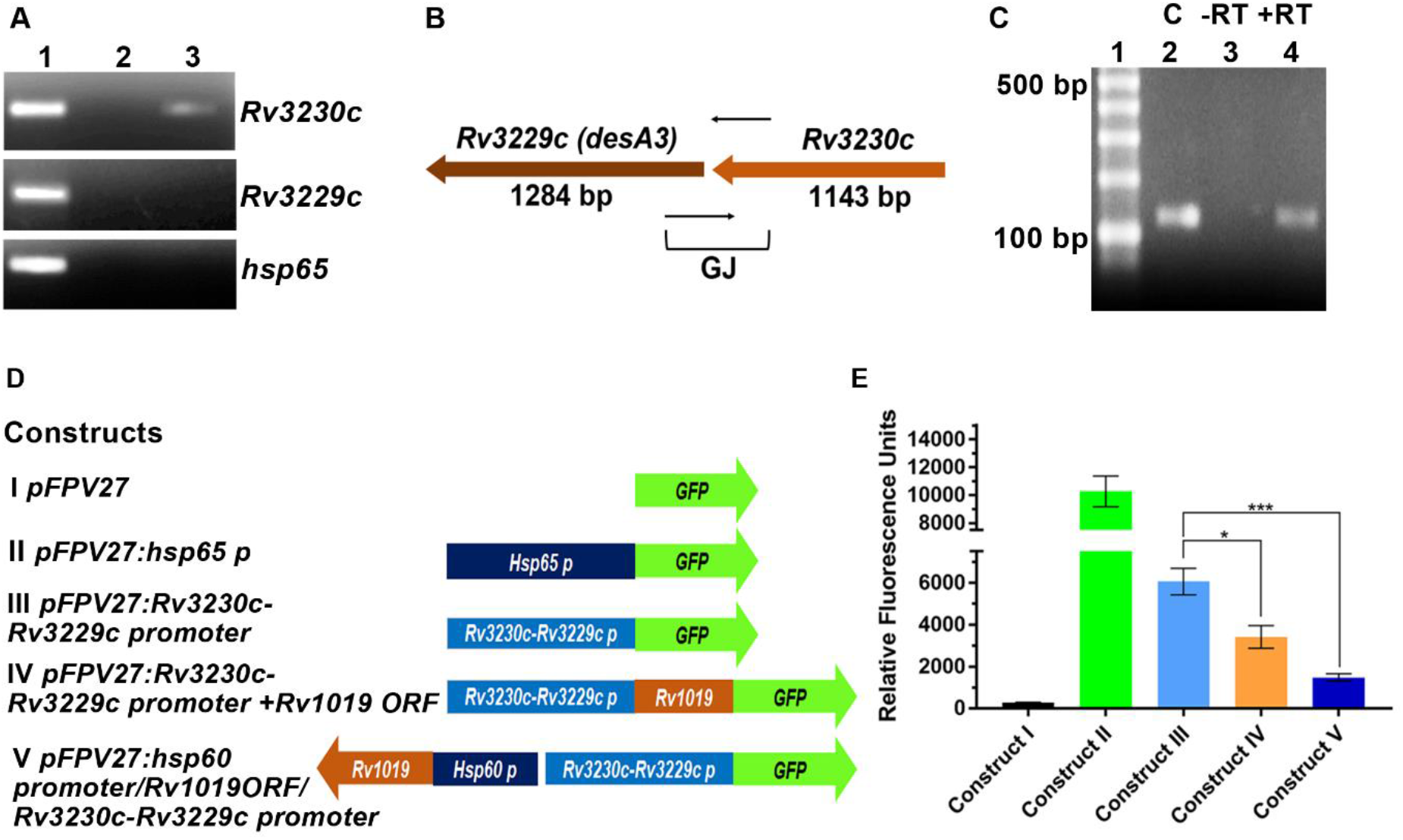
*Rv3230* and *Rv3229* have a common promoter. **(A)** ChIP assay to demonstrate *in vivo* binding of Rv1019 to *Rv3230c-Rv3229c* promoter. Sheared *M. tuberculosis* DNA– protein complex was immunoprecipitated with antibodies against Rv1019, and the DNA in the immune complexes was analyzed by PCR using specific primers for *Rv3230c, Rv3229c* and *hsp65* promoters. Lane 1: PCR amplicon of input DNA, Lane 2: PCR amplicon of IgG pull-down, Lane 3: PCR amplicon of Rv1019 pull-down; (**B**) Orientation and size of *Rv3230c* and *Rv3229c*. Gene junctions are indicated (GJ). Thin black arrows represent the DNA region selected to design PCR primers to amplify the gene junctions; (**C**) PCR amplification to prove cotranscription of *Rv3230c* and *Rv3229c*. Lane 1: 100 bp DNA ladder, Lane 2: positive control, Lane 3: PCR amplification from negative reverse transcription reaction, Lane 4: PCR amplification from positive reverse transcription reaction; (**D**) Reporter constructs. Construct I: promoter-less *pFPV27* vector (negative control); construct II: vector carrying *hsp65* promoter (positive control); construct III: -1 to -300 region upstream of *Rv3230c* cloned upstream of GFP. Construct IV: *Rv1019* ORF cloned upstream of GFP downstream to *Rv3230c* promoter as a transcriptional fusion. Construct V: *Rv1019* under *hsp60* promoter in reverse orientation of GFP; (**E**) Reporter assay in *M. smegmatis*. Each of the afore-mentioned constructs was electroporated into *M. smegmatis ΔMSMEG_5424* strain, and the fluorescence was measured after 48 h and represented in terms of RFU. GFP expression from construct III vs construct IV, and construct III vs construct V were compared, and values are represented as mean ± SD of three independent experiments (Tukey’s multiple comparisons test). ***P ≤ 0.0003 and *P ≤ 0.01.

*In silico* operon prediction tools and expression arrays showed that *Rv3230c* and its gene neighbourhood *Rv3229c* (*desA3*) could be cotranscribed [16] [17]. These two proteins are involved in the synthesis of oleoyl-CoA from stearoyl-CoA. To check whether these two genes are cotranscribed, we designed PCR primers that can amplify the junctions between *Rv3230c* and *Rv3229c* (**Fig. 1B**). By performing PCR on cDNA of normally grown *M. tuberculosis*, we were able to amplify the *Rv3230c-Rv3229c* junction (**Fig. 1C**). Similarly, we checked if the homologues in *M. smegmatis MSMEG_1885* (homolog of *Rv3230c*) and *MSMEG_1886* (homolog of *Rv3229c*) are cotranscribed, and we were able to amplify *MSMEG_1885*-*MSMEG_1886* junction (**Fig. S1 A** and **B**). These results indicate that *Rv3230c* and *Rv3229c* are transcribed together into a single mRNA and Rv1019 could have a common binding site upstream of *Rv3230c*.

To analyse the regulatory activity of Rv1019 on this promoter region, we created a series of GFP reporter constructs in pFPV27 reporter system (**Fig. 1D; S2 A** and **B**). All the constructs were then electroporated into the *M. smegmatis ΔMSMEG_5424* strain to avoid any non-physiological artefacts associated with the coexpression of *MSMEG_5424* and *Rv1019* in the same host. GFP expression from the strains harbouring these constructs were monitored. By measuring GFP, we found that *Rv3230c-Rv3229c* promoter is active in *M. smegmatis*, and the strain harbouring promoter with *Rv1019* ORF (construct IV) showed significant reduction in GFP expression compared to bacteria carrying construct III (plasmid containing only the *Rv1019* promoter; **Fig. 1E**). For further validation, *Rv3230c-Rv3229c* promoter was cloned upstream of GFP, and into the same construct we subcloned *Rv1019* ORF under the constitutive *hsp60* promoter in the opposite orientation (construct V; **Fig. 1D**). Interestingly, *M. smegmatis* with these constructs showed a significant reduction in green fluorescence when compared to that carrying the promoter-GFP alone controls (**Fig. 1E**). These results confirmed that Rv1019 can bind to a single promoter region upstream of *Rv3230c-Rv3229c* and repress their expression. This was further confirmed by analysing GFP expression by confocal photomicrography (**Fig. S2 C**).

### Rv1019 binds to a 12-bp inverted repeat on *Rv3230c-Rv3229c* promoter

To identify the binding sequence of Rv1019 on *Rv3230c-Rv3229c* promoter, we performed an EMSA using the 300-bp upstream sequence (5 nM) with increasing concentrations of recombinant Rv1019 (2–10 µM). Formation of a distinct DNA–protein complex was observed intensity of which increased as a function of the concentration of the protein. Specificity of binding was proven by competitive and noncompetitive EMSA (**Fig. 2A** and **2B**). In competitive EMSA, 10-fold specific template was used. In contrast, binding was not observed even at 10-fold excess of nonspecific promoter (of an irrelevant *M. tuberculosis* gene *Rv0440c* that codes for GroEL2) confirming that binding is specific.

**Fig. 2.**
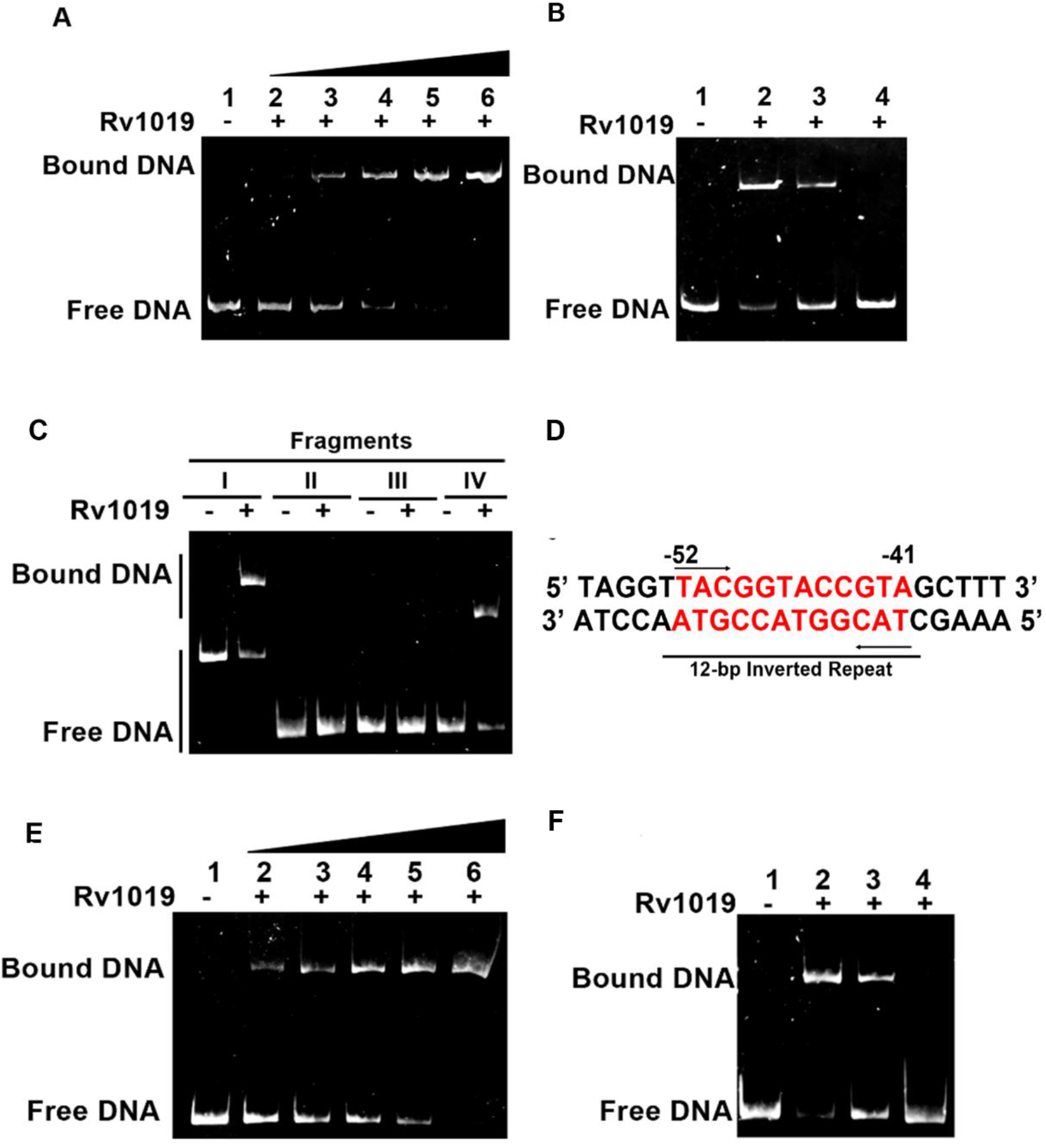
Rv1019 binds to a 12-bp inverted repeat on *Rv3230c-Rv3229c* promoter. **(A)** EMSA shows *in vitro* binding of Rv1019 to -1 to -300 region upstream of *Rv3230c* ORF. Lane 1: DNA without Rv1019, Lanes 2–6: increasing concentrations of Rv1019 (2–10 μM); **(B)** Competitive and noncompetitive EMSA to validate the specificity of binding. Lane 1: DNA without protein, Lane 2: DNA incubated with 10 μM of Rv1019 (positive control), Lane 3: competitive EMSA, Lane 4: noncompetitive EMSA (non-specific template incubated in the presence of 10 μM of Rv1019); (**C**) EMSA with DNA fragments upstream of ORF. Lanes 1–2: -1to -300 upstream, Lanes 3–4: -200 to -300 upstream, Lanes 5–6: -100 to -200 upstream, Lanes 7–8: -1 to -100 upstream; (**D**) 12-bp inverted repeats identified in -1 to - 100region; (**E**) EMSA on 12-bp palindrome. Lane 1: palindromic DNA sequence without Rv1019, Lanes 2–6: DNA with increasing concentrations of Rv1019; **(F)** Competitive and noncompetitive EMSA to validate the binding of Rv1019 on the palindromic sequence. Lane 1: DNA without protein, Lane 2: with DNA incubated with 10 μM of Rv1019 (positive control), Lane 3: competitive EMSA, Lane 4: non-competitive EMSA with a nonspecific DNA.

To determine the exact binding site of Rv1019 on *Rv3230c-Rv3229c* promoter, PCR-amplified fragments of different lengths were subjected to EMSA after incubating them with the recombinant protein (**Fig. S3 A**). At 10 µM concentration of recombinant Rv1019, we observed formation of DNA–protein complexes as evidenced by a distinct shift in the migration pattern of DNA fragments I (−1 to -300) and IV (−1 to -100), while the fragments II (−200 to -300) and III (−100 to -200) did not show any shift in the presence of the protein **(Fig. 2C**). These results suggested that the binding region is located between -1 and -100 from the start site of the *Rv3230c. In silico* analysis of the -1 to -100 of the upstream region of *Rv3230c* ORF revealed a 12-bp perfect inverted repeat sequence (**Fig. 2D**). EMSA using this region as the template showed formation of a DNA–protein complex and the binding on this was further validated with competitive and noncompetitive EMSA (**Fig. 2E** and **2F**). This was further confirmed *in vivo* employing a reporter construct (construct VI) containing only the -100 to -300 region (without the binding sequence) (**Fig. S3 B**). Increased fluorescence was observed in *M. smegmatis* carrying the construct VI in comparison with the strain carrying the construct IV which contained the cognate binding site (**Fig. S3 C** and **D**). These results confirmed that the Rv1019 binds to the 12-bp inverted repeats upstream of *Rv3230c* leading to the repression of *Rv3230c-Rv3229c* promoter activity *in vivo*. Similarly, analysis of the upstream region of *MSMEG_1885* (homologue of *Rv3230c*) in *M. smegmatis* revealed the same 12-bp inverted repeat at -7 to -18 bp region. This confirmed that this region is conserved in *M. tuberculosis* and *M. smegmatis* (**Fig. S4 A**). Binding of Rv1019 to this region was confirmed by performing an EMSA (**Fig. S4 B** and **C**).

### Rv1019 is upregulated in Wayne’s hypoxia-induced dormancy

Previously, to analyse the proteome, we induced dormancy in *M. tuberculosis* by employing Wayne’s hypoxia-induced dormancy model [8]. We used this method to validate the expression of *Rv1019*, and the self-generated hypoxia was monitored using methylene blue indicator (**Fig. 3A**). The initial bluish green colour gradually became colourless, to disappear completely on day eight which indicated attainment of microaerophilic condition inside the tube. The non-replicating persistent stage 1 (NRP1) was attained on the 12^th^ day and the non-replicating persistent stage 2 (NRP2) or enduring hypoxia was attained on the 21^st^ day as were observed by others using Wayne’s model [18]. NRP2 mimics the *in vivo* situation where *M. tuberculosis* attains dormancy inside the macrophages. Viability of *M. tuberculosis* subjected to aerobic growth and hypoxia were checked by growth curve analysis to confirm induction of dormancy (**Fig. 3B**). Bacteria were harvested from normal aerobic culture, NRP1, NRP2 stages and were subjected to Quantitative Real-Time PCR and Western blotting to validate the expression of *Rv1019*. Expression of *Rv1019* at various stages of dormancy showed differential expression of the gene (**Fig. 3C**). *Rv1019* expression was found to be significantly upregulated during NRP2 stage compared to that under normal aerobic conditions. Expression of *devR*, a dormancy regulatory protein was used as the positive control and *sigA* as an endogenous control. Expression of *Rv1019* during dormancy was analysed by western blotting as well with Rv1019 antibodies (**Fig. 3D**). Thus, we confirmed that *Rv1019* is differentially expressed with upregulation in dormancy (NRP2) compared to aerobic growth.

**Fig. 3.**
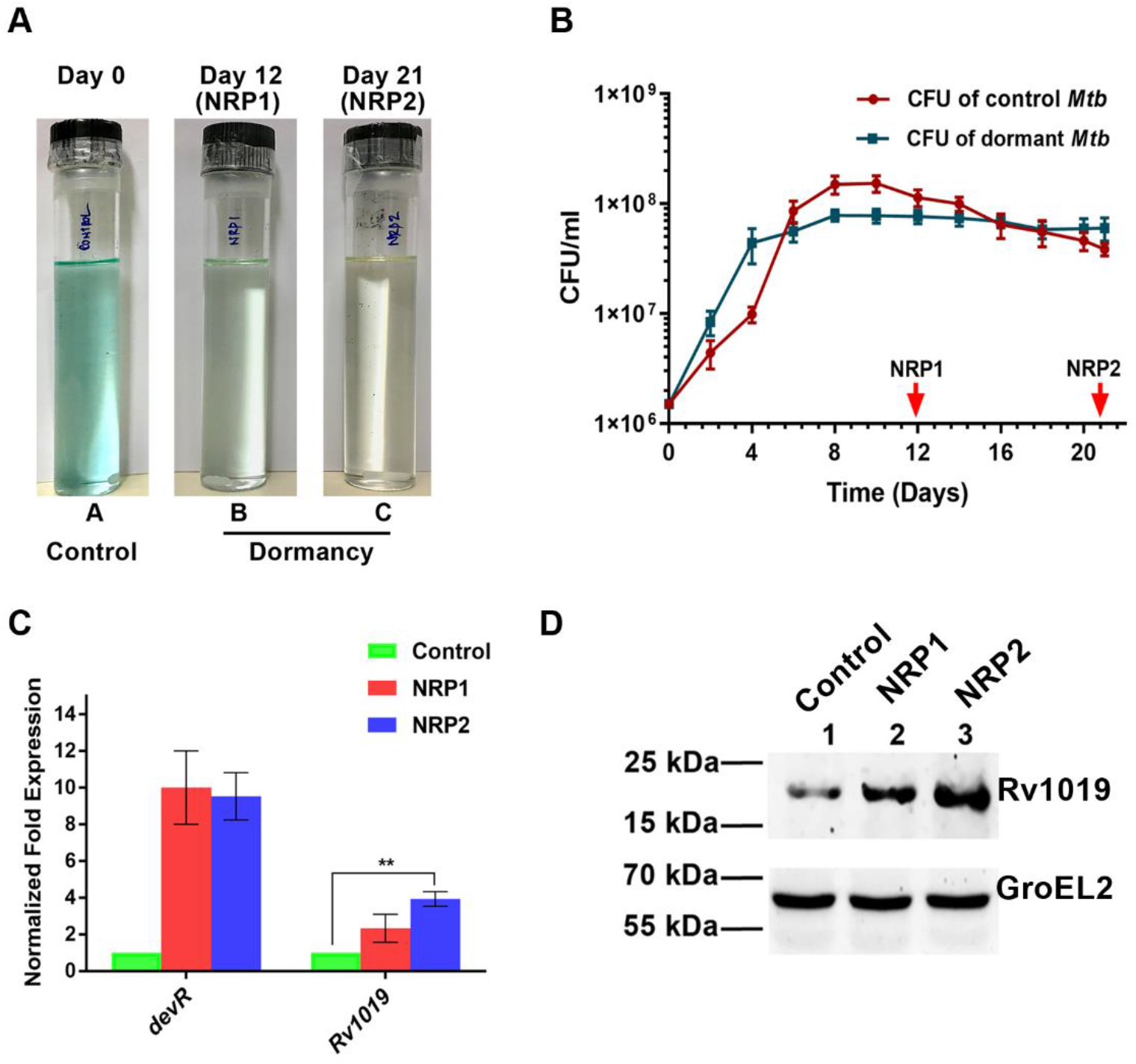
Wayne’s hypoxia-induced dormancy model of *M. tuberculosis* and analysis of *Rv1019* expression. **(A)** Experimental setup for dormancy shows the control tubes to monitor the levels of oxygen. All the tubes contain methylene blue indicator: Tube A represents control (first day); Tubes B & C contain dormant *M. tuberculosis* incubated for 12 (NRP1) and 21 (NRP2) days, respectively (methylene blue loses its colour); (**B**) Measurement of growth of *M. tuberculosis* H37Rv by optical density and CFU (values at each point of time represent an average value from three tubes). CFU values of *M. tuberculosis* subjected to dormancy and normal aerobic growth. Red arrows indicate sampling points; (**C**) Quantitative Real-Time PCR analysis. Relative fold expression values of *Rv1019* and *devR* at various stages of dormancy and reactivation. Values are the mean ± standard deviation of three independent experiments (Student’s *t*-test). **P ≤ 0.001; (**D**) Western blotting showing the expression of *Rv1019* during dormancy and reactivation. Lane 1: lysate of aerobically grown *M. tuberculosis*, Lane 2: lysate of *M. tuberculosis* from the 12^th^ day of dormancy (NRP1), Lane 3: lysate of *M. tuberculosis* from the 21^st^ day of dormancy (NRP2).

### Rv1019 is recruited to *Rv3230c-Rv3229c* promoter in hypoxia

To analyse if Rv1019 is recruited to *Rv3230c-Rv3229c* promoter during hypoxia, we performed chromatin immunoprecipitation followed by PCR. We found a very high enrichment of Rv1019 on *Rv3230c-Rv3229c* promoter, as evidenced by the thick band of the amplicon, during NRP2 compared to normal aerobic growth (**Fig. 4A**). This confirmed that Rv1019 is recruited to *Rv3230c*-*Rv3229c* promoter during hypoxia. Interestingly, we also found that Rv1019 does not show binding on its own promoter during NRP2. At the same time, we checked the expression of *Rv3230c* and *Rv3229c* by qRT-PCR analysis in hypoxia and found that *Rv3230c* and *Rv3229c* are significantly downregulated during hypoxia compared to normal aerobic conditions of growth (**Fig. 4B**). To analyse if the overexpression of *Rv1019* could cause downregulation of these genes, we used *M. tuberculosis* overexpressing *Rv1019*. Overexpression of *Rv3230c* and *R3229c* was analysed by qRT-PCR and was confirmed by SDS-PAGE and western blotting (**Fig. S5 A** and **B**). We found that *M. tuberculosis* constitutively expressing *Rv1019* showed significant downregulation of expression of *Rv3230c* and *Rv3229c* (**Fig. 4C**). Also, we analysed the expression of *MSMEG_1885* and *MSMEG_1886*, in *M. smegmatis* constitutively expressing *Rv1019* and *M. smegmatis* constitutively expressing *MSMEG_5424* (*Rv1019* homolog). We found significant downregulation of *MSMEG_1885* and *MSMEG_1886* (**Fig. S6 A** and **Fig. S6 B**). This confirmed that overexpression of *Rv1019* negatively regulates *Rv3230c-Rv3229c* operon in *M. tuberculosis*, and overexpression of *MSMEG_5424* negatively regulates *MSMEG_1885-MSMEG_1886* operon in *M. smegmatis*. Interestingly, we did not observe any change in the expression of *MSMEG_1885* and *MSMEG_1886* in the case of *M. smegmatis* deficient in *MSMEG_5424* (**Fig. S6 C**).

**Fig. 4.**
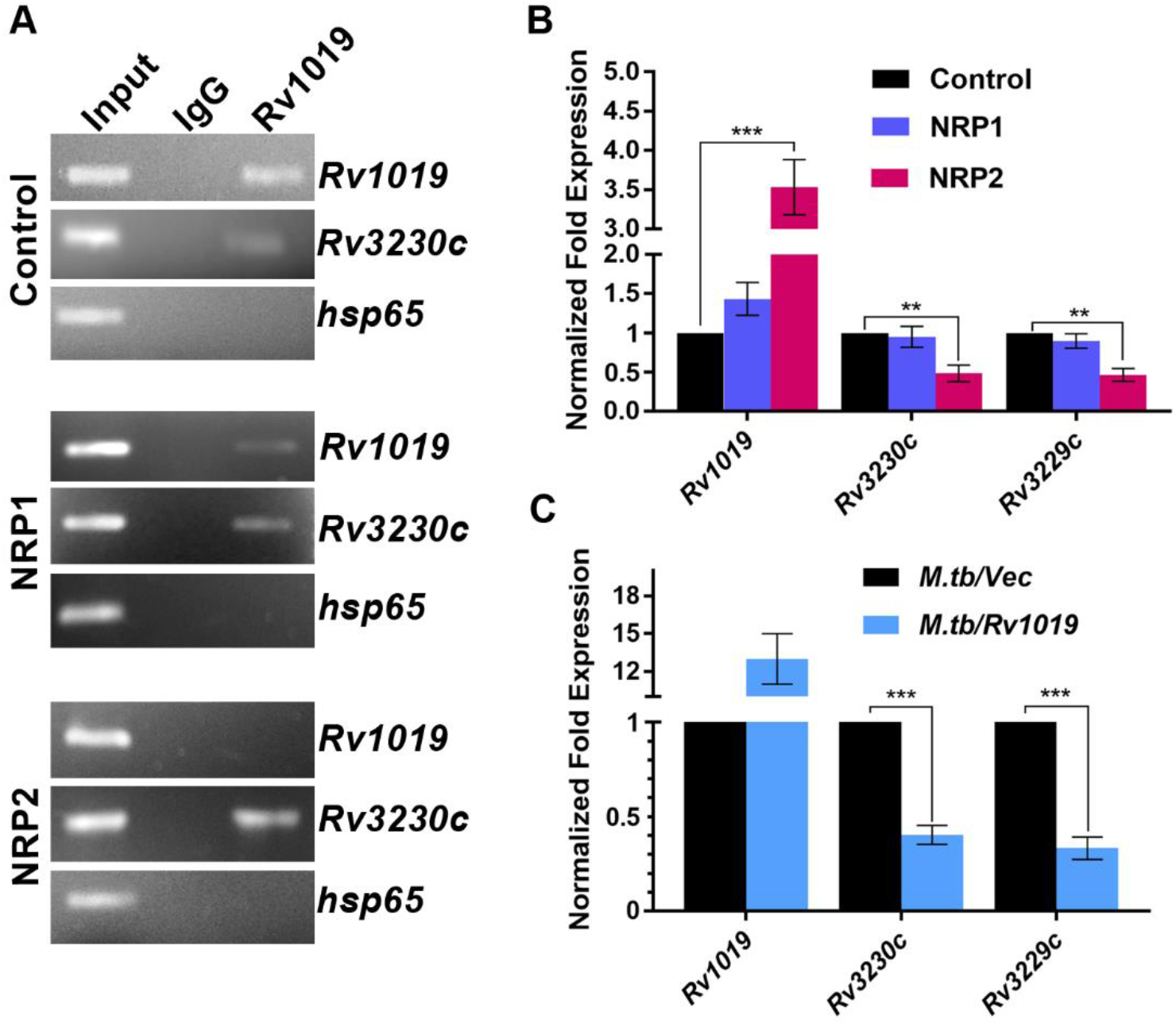
Rv1019 is recruited to the *Rv3230c-Rv3229c* promoter and downregulates the operon in *M. tuberculosis*. **(A)** ChIP PCR showing binding of Rv1019 on its own promoter and that of *Rv3230c-Rv3229c* during dormancy compared to normal aerobic growth (control); (**B**) Quantitative real-time PCR of *Rv1019, Rv3230c and Rv3229c* in dormancy compared to normal aerobic growth; **(C)** Quantitative Real-Time PCR showing expression of *Rv1019* (constitutive), and the expression of endogenous *Rv3230c* and *Rv3229c*. Values are the mean ± standard deviation of three independent experiments (Student’s *t*-test) ***P ≤ 0.001, **P ≤ 0.01.

### Rv1019 negatively regulates *Rv3230c-Rv3229c* operon in hypoxia in *M. smegmatis*

To validate the expression of *Rv3230c-Rv3229c* in hypoxia, we used *M. smegmatis* as a model organism. First, we induced dormancy in wild type and *ΔMSMEG_5424* strains of *M. smegmatis*. Induction of dormancy in *M. smegmatis* was confirmed by colour change in methylene blue (**Fig. 5A**). Unlike *M. tuberculosis, M. smegmatis* attains NRP1 on the 7^th^ day and NRP2 on the 10^th^ day. Viability and growth of the wild type and the deletion mutant strains during induction of dormancy were determined by counting CFU and by measuring the turbidity of the culture at 600 nm. Both strains showed similar growth pattern during hypoxia (**Fig. S7 A**).

**Fig. 5.**
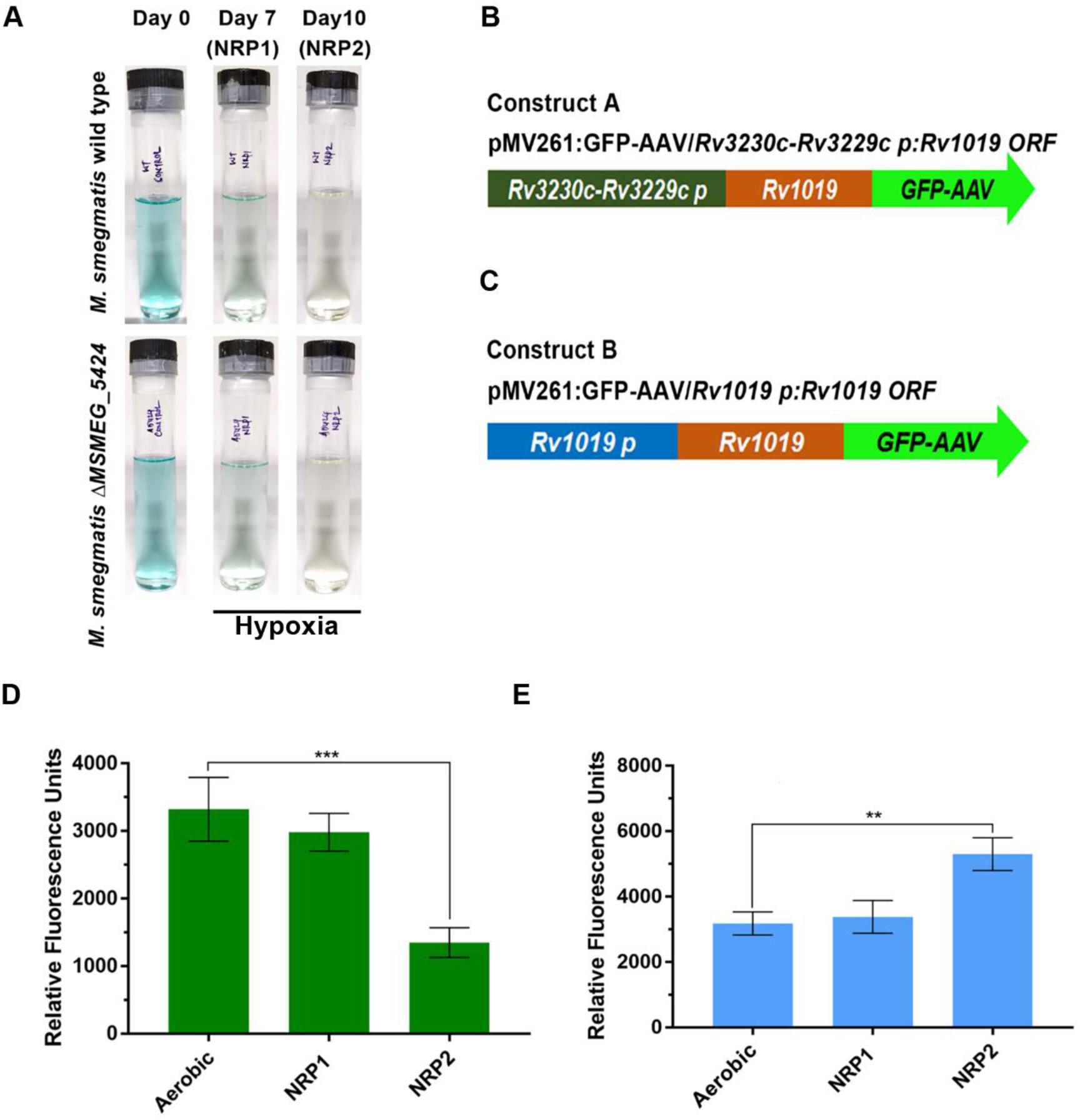
Induction of dormancy in *M. smegmatis*, and reporter assay. **(A)** Experimental setup for dormancy to monitor the levels of oxygen: all the tubes contain methylene blue indicator: Tube A represents day 0; tubes B & C contain dormant *M. smegmatis* incubated for 7 (NRP1) and 10 (NRP2) days, respectively; (**B**) Schematic diagram of construct A; (**C**) Schematic diagram of construct B; (**D**) Reporter assay of *M. smegmatis* harbouring construct A. Values are the mean ± SD of three independent experiments (Tukey’s multiple comparisons test). ***P ≤ 0.001; (**E**) Reporter assay of *M. smegmatis* harbouring construct B. Values are the mean ± SD of three independent experiments (Tukey’s multiple comparisons test). **P ≤ 0.003.

To analyse the *Rv3230c-Rv3229c* promoter and *Rv1019* promoter activity under hypoxia, we generated reporter constructs with pMV261:*GFP-AAV* as the backbone vector (**Fig. S7 B**). GFP-AAV is a GFP protein with a short half-life (∼2 h) and provides an accurate representation of promoter activity in real-time. We have generated a reporter construct in which we cloned *Rv1019* ORF upstream of *GFP-AAV*. To this construct, we cloned *Rv3230c-Rv3229c* promoter (containing the 12-bp cognate binding site) upstream to *Rv1019 ORF* (Construct A: pMV261:*GFP-AAV*/*Rv3230c-Rv3229c* promoter/*Rv1019 ORF*; (**Fig. 5B** and **Fig. S7 C**). Similarly, we made another construct in which we cloned *Rv1019* ORF upstream of *GFP-AAV Rv1019* promoter. This construct was further modified by cloning *Rv1019* promoter upstream of *Rv1019 ORF* (Pushparajan *et al*., 2020) (Construct B: pMV261:*GFP-AAV*/*Rv1019* promoter/*Rv1019 ORF*; (**Fig. 5C** and **Fig. S7 D**). These constructs were then transformed individually into *M. smegmatis ΔMSMEG_5424* to avoid potential non-physiological artefacts associated with the expression of both *Rv1019* and *MSMEG_5424* in the same strain.

Dormancy was induced in *M. smegmatis ΔMSMEG_5424* harbouring Construct A and *M. smegmatis ΔMSMEG_5424* harbouring Construct B. GFP expression during NRP1 and NRP2 stages was compared with that in aerobically grown bacteria, and the values were represented in terms of RFU. We found significant reduction in fluorescence during NRP2 stage in *M. smegmatis ΔMSMEG_5424* harbouring Construct A (**Fig. 5D**), whereas *M. smegmatis ΔMSMEG_5424* harbouring Construct B showed significant increase in fluorescence (**Fig. 5E**). This was further confirmed by confocal photomicrography (**Fig. S8 E** and **F**). These results confirmed that compared to aerobically grown bacteria, Rv1019 promoter activity is significantly high in dormant bacteria where Rv1019 is recruited to the promoter of *Rv3230c-Rv3229c* operon and represses its expression.

## Discussion

Success of *M. tuberculosis* as an intracellular pathogen depends on its ability to overcome the challenging environment in the host, and the bacterium continues to be the leading cause of death by an infectious agent. The main obstacle in eradicating tuberculosis, in spite of effective drugs and the BCG vaccine, is the ability of *M. tuberculosis* to persist in macrophages in a dormant form and get reactivated when the immune system of the host becomes weak. *M. tuberculosis* has approximately 4000 genes coding for proteins [5] [19] of which about 40% is still uncharacterized and fall under the category of ‘hypothetical’ proteins.

Transcriptional regulation in intracellular pathogens plays an important role in their adapting to the challenging environment inside the host cells. Transcriptional regulators are key proteins in any biological system as they regulate all biological functions in a cell. *M. tuberculosis* has 214 TRs [12] and many of them are still hypothetical. Characterization of each of them is critical to unravel the downstream biochemical processes controlled by them. This will enable us to devise better and more effective strategies to combat tuberculosis.

Most of the transcriptional regulators in *M. tuberculosis* are multifunctional in nature [8]. Chromatin immunoprecipitation followed by microarray hybridization or high-throughput sequencing (ChIP-chip and ChIP-seq) studies have been applied to study genome-wide gene regulation in *M. tuberculosis* and this helped to map the transcriptional regulator binding sites [15] [12]. These techniques have helped to map the transcriptional regulatory network of *M. tuberculosis* for a better understanding of the adaptive mechanism employed by the pathogen to survive in the host.

In our previous study, we had shown that under normal aerobic conditions of growth Rv1019 acts as an autorepressor and regulates its own expression [13]. Also, we showed that *Rv1019* and its downstream genes *Rv1020* and *Rv1021* are cotranscribed and Rv1019 negatively regulate this operon. However, on searching the transcription factor overexpression databases and the transcriptional regulatory networks of *Mtb* [12] we found that Rv1019 could be involved in regulating other genes as well.

In the present study, we investigated the role of Rv1019 in regulating the expression of *Rv3229c* (*desA3*) and *Rv3230c* in *Mtb*. The proteins coded by these genes are involved in the biosynthesis of oleoyl-CoA, a precursor of membrane phospholipids [20] [21]. We validated the recruitment of Rv1019 to *Rv3230c*and *Rv3229c* promoter by ChIP assay followed by PCR and found that Rv1019 can bind upstream of *Rv3230c* ORF but not of *Rv3229c*. This indicated that *Rv3230c* and *Rv3229c* could be adjacent genes under a single promoter which is upstream of *Rv3230c*. Operon prediction tools and expression arrays showed that *Rv3230c* could be part of an operon with its downstream gene *Rv3229c* [16] [17] [22]. We have validated this by RT-PCR using primers which can amplify the junction between *Rv3230c* and *Rv3229c*. Our results agree with the previous studies which explained the functional relationship between these proteins [21]. Similarly, the homologues *MSMEG_1885 (∼Rv3230c*) and *MSMEG_5424 (∼Rv3229c*) in *M. smegmatis* were also found to be cotranscribed. Thus in *M. tuberculosis Rv3230c* and *Rv3229c* form a two gene operon in which *Rv3230c* is the first gene, and the *Rv3230c-Rv3229c* locus is conserved in *M. tuberculosis* and *M. smegmatis*. Also, from the mycobacterium data bases we found that this locus is conserved in other mycobacterium species such as *M. bovis, M. leprae* and *M. marinum*. Conservation of this operon in both pathogenic and non-pathogenic strains of mycobacterium indicates that the role of these proteins in growth of the bacteria is irrelevant to pathogenicity.

Employing reporter assays we showed that Rv1019 is a negative regulator of *Rv3230c-Rv3229c* expression. Mapping of the cognate binding site revealed a 12 bp perfect inverted repeat upstream (−41 bp to -52 bp) of the *Rv3230c* ORF. We found that this sequence is present within the Rv1019 binding region upstream of *Rv3230c* as demonstrated in a previous study using transcriptional regulator overexpression followed by ChIP-seq [12]. Interestingly, we found there is no homology at all between *Rv1019* and the *Rv3230c-Rv3229c* promoters. Thus, this protein has the ability to bind to two different DNA sequences which is a characteristic feature of TetR family of TRs [23] [24]. We also found that the 12-bp cognate binding site is conserved in *M. smegmatis* upstream of *MSMEG_1885*, the homologue of *Rv3230c*.

Proteomic profiling studies have shown that *Rv1019* is significantly upregulated in Wayne’s hypoxia-induced dormancy model [8] [14]. Employing this model, we induced non-replicating persistence in *Mtb*. Further, using quantitative real-time PCR analysis we checked the expression of *Rv1019. DevR*, a well characterised transcriptional regulator of *M. tuberculosis* which is upregulated during dormancy and reactivation [25] [8] was used as a positive control. Our results show that *Rv1019* is differentially expressed during dormancy and reactivation with a significant upregulation during NRP2 stage of dormancy. This was further validated by western blotting *M. tuberculosis* lysates from dormancy and reactivation using antibodies against Rv1019. Our results agree with the previous studies where *Rv1019* was found to be differentially expressed and showed similar profile [8] [14].

In *M. tuberculosis, Rv3230c* and *Rv3229c* code for enzymes involved in the biosynthesis of oleic acid from steric acid. Oleic acid, an unsaturated 18-carbon fatty acid, is a major precursor of mycobacterial membrane phospholipids. Rv3229c (DesA3) forms a functional complex with the corresponding oxidoreductase Rv3230c and converts saturated stearic acid into unsaturated oleic acid in the presence of molecular oxygen and NADPH [21] [26].

Deletion of *MSMEG_1886* (*desA3*) was partially auxotrophic for oleic acid [27]. Oleic acid synthesis is essential for the cell wall formation of actively replicating bacteria. To adapt to the intracellular stress conditions during dormancy, *M. tuberculosis* has developed strategies to regulate several pathways and downregulates genes involved in the synthesis of several membrane lipids [28] [29].

To validate if Rv1019 is recruited to *Rv3230c*-*Rv3229c* promoter under dormancy, ChIP assay followed by PCR was performed. We found significant enrichment of *Rv3230c-Rv3229c* promoter in NRP2 compared to the control. This was further confirmed by reporter assays using *M. smegmatis*, where we found very low fluorescence in bacteria harbouring reporter construct with *Rv3230c-Rv3229c* promoter whereas constructs harbouring *Rv1019* promoter showed significant increase of fluorescence. Based on these observations we propose that *Rv1019* is upregulated under dormancy and the protein is recruited to the promoter of *Rv3230c-Rv3229c* and downregulates this operon.

Even though we found binding of Rv1019 on *Rv3230c-Rv3229c* promoter during normal growth, it had very less influence on the expression of *Rv3230c* and *Rv3229c*. As evident form our previous study, we believe that under normal aerobic growth Rv1019 acts as an autorepressor and majority of the protein is bound to its own promoter [13]. However, during upregulation/overexpression, Rv1019 was found to have increased affinity on *Rv3230c-Rv3229c* promoter. This was evident from our reporter assays and ChIP-PCR where we found enhanced recruitment of Rv1019 to *Rv3230c* promoter under hypoxia-induced dormancy compared to normal growth.

Further, quantitative real-time PCR showed significant downregulation of *Rv3230c* and *Rv3229c* during NRP2 compared to that under normal growth. Constitutive expression of *Rv1019* in aerobic conditions leads to the downregulation of *Rv3230c* and *Rv3229c* in *M. tuberculosis* and its homologues *MSMEG_1886* and *MSMEG_1885*, respectively, in *M. smegmatis*. This confirmed that Rv1019 can negatively regulate *Rv3230c-Rv3229c* operon. This is in accordance with the previous study where vitamin C-induced dormancy model showed significant downregulation of *Rv3229c* (*desA3*) [30]. At the same time, expression of *MSMEG_1885* (≈*Rv3230c*) and *MSMEG_1886* (≈*Rv3229c*) in *ΔMSMEG_5424* mutant of *M. smegmatis* did not show any difference in expression compared to the wild-type strain. This, we presume, could be due to poor recruitment of *MSMEG_5424* (≈*Rv1019*) on this promoter or the promoter is controlled by a different TR with higher affinity under aerobic growth.

Interestingly, even though Rv1019 is an autorepressor, we did not observe binding of Rv1019 to its own promoter during hypoxia. At this point we are unclear about the loss of affinity of Rv1019 for its own promoter during hypoxia. Transcriptional regulator overexpression databases shows that regulators such as Rv1033c, Rv3574, Rv0023 and Rv3133c can bind to *Rv1019* promoter [15]. Upregulation of *Rv1019* during dormancy, could probably be result of positive regulation by any of this or a different regulator with high affinity for *Rv1019* promoter; however, more studies are required to validate this hypothesis.

Even though this study revealed the role of Rv1019 in regulating the expression of *Rv3230c-Rv3229c* operon in *M. tuberculosis*, we are not sure of the other roles played by Rv1019 during hypoxia. Turkarslan *et al*., 2015 showed that, apart from binding to *Rv3230c* promoter, Rv1019 can bind to *Rv3573c* and *Rv3574* promoters as well which are involved in cholesterol metabolism in *M. tuberculosis. Rv3574* (*kstR*) controls the expression of several genes involved in cholesterol catabolism [31] [32]. Role of Rv1019 in regulating the expression of these genes must be further explored. After all, 214 TRs regulate the expression of about 4000 genes in *M. tuberculosis* which suggests that each TR could regulate about 20 genes. Considering these facts, we hypothesize that Rv1019 may have a vital regulatory role in controlling the expression of genes involved in lipid metabolism during dormancy.

## Methods

### Bacterial strains, plasmids, and growth conditions

All *E. coli* strains used in this study were grown in Luria Bertani (LB) broth or on LB agar (HiMedia, Mumbai, India) at 37 °C. All steps involving handling of *Mtb* were carried out in a biosafety level three (BSL3) facility. *M. tuberculosis* and *M. smegmatis* were cultured in Middlebrook 7H9 (Becton Dickinson, Franklin Lakes, NJ, and USA) or Middlebrook 7H10 supplemented with 10% albumin-dextrose-catalase (HiMedia, Mumbai, India). For promoter analysis in *M. smegmatis* pFPV27 (Addgene), pMV261 (Krishna Kurthkoti) vectors were used and for overexpression studies pBEN (Lalitha Ramakrishnan) vector was used (**Table 1**). PCR primers and oligos were purchased from Sigma-Aldrich (**Table 2)**. Restriction enzymes were purchased from New England Biolabs, Ipswich, MA, USA. All other chemicals and reagents were purchased from Sigma-Aldrich, St. Louis, MO, USA or USB, Cleveland, OH, USA.

### Chromatin immunoprecipitation (ChIP) assay

ChIP assay was performed according to previously described protocols [33]. Mid-log phase *M. tuberculosis* or *M. smegmatis* culture (A_600_ ∼0.6) was treated with 1% formaldehyde, and incubated at 25 °C for 30 mins, to facilitate DNA-protein cross-linking, and was terminated by addition of glycine to a final concentration of 0.5 M at 25 °C and incubating for 10 mins. After washing in phosphate buffered saline (PBS; pH 7.4) containing 1 mM phenylmethylsulfonyl fluoride (PMSF), the cells were centrifuged at 3000g for 10 min at 4°C, pellets were resuspended in ChIP lysis buffer [50 mM HEPES-KOH (pH 7.5), 150 mM NaCl, 1 mM EDTA, 1% Triton X-100, 0.1% (w/v) sodium deoxycholate, 0.1% SDS, 0.1 mg/mL RNaseA, 1X protease inhibitor cocktail and 1 mM PMSF]. Cells were lysed in a Mini Bead Beater (Biospec, Bartlesville, OK, USA) by 3 pulses, 30 sec each at 42 × 100 rpm, using zirconium beads (0.1 mm). DNA was sheared using a Bioruptor sonicator (Diagenode, Seraing, Belgium) with 35 cycles of 30 sec on/off pulses to generate fragments of ∼500 bp. DNA fragments were recovered by centrifugation at 12000 g for 10 mins at 4 °C. To compare the enrichments in various conditions, total protein concentration of the lysate was estimated and equal quantity of protein from each condition was used for shearing. One hundred microlitre of this lysate was used as the input control for ChIP. The lysate was pre-cleared by incubating with 10 μL of magnetic beads (Diagenode) on a rotary shaker for 45 mins at room temperature. The supernatant was removed and incubated with either pre-immune serum or antibodies against Rv1019 and incubated on a rotary shaker overnight at 4 °C. The samples were then incubated with 30 μL of pre-blocked magnetic beads for 30 mins at 4 °C. After incubation, samples were washed with ChIP lysis buffer, followed by four washes each with ChIP lysis buffer containing 500 mM NaCl, wash buffer [10 mM Tris (pH 8.0), 250 mM LiCl, 1 mM EDTA, 0.5% Igepal CA-630 and 0.5% sodium deoxycholate] and TE buffer (pH 7.5). Immunocomplexes were eluted from the magnetic beads in 100 μL elution buffer [10 mM Tris (pH 7.5), 10 mM EDTA and 1% SDS] at 65 °C for 30 mins. Both immunoprecipitated and input DNA were subjected to Proteinase K treatment (80 μg/100 μL). DNA was extracted using phenol-chloroform (1:1) and precipitated using 100% ethanol and dissolved in 10 mM Tris (pH 8.0). PCR was carried out using specific primers.

### Generation of reporter constructs

Reporter constructs were made as previously described [13] [34]. To study the promoter activity of *Rv3230c-Rv3229c*, a 300 bp DNA fragment upstream from the start site of *Rv3230c* was amplified from *M. tuberculosis* H37Rv genomic DNA, and cloned into pFPV27, using restriction sites *Bam*HI and *Eco*RI, upstream of GFP. To analyse the effect of Rv1019 on *Rv3230c-Rv3229c* promoter, another construct was made where 300 bp DNA fragment upstream from the start site of *Rv3230c* was cloned into pFPV27, using restriction sites *Bam*HI and *Eco*RI, and *Rv1019* ORF was cloned downstream at *Eco*RI and *Kpn*I sites upstream of GFP.

To confirm the regulation of *Rv3230c-Rv3229c* by *Rv1019, Rv1019* ORF was first cloned into pBEN vector, downstream of constitutive promoter *hsp60* using *Hin*dIII and *Bam*HI sites. The *hsp60-Rv1019* region was PCR amplified and cloned into pFPV27 in reverse orientation with respect to *GFP* to avoid any read through effect of *hsp60* on *GFP* expression. Transcriptional terminator T4g32 was then cloned downstream of *Rv1019*. The promoter of *Rv3230c-Rv3229c* was cloned upstream of *GFP* into *Apa*I site of the modified pFPV27 in the same orientation as *GFP*. Constructs were transformed into *M. smegmatis* and the transformants were selected on 7H10 agar plates containing kanamycin (50 μg/mL). The recombinant *M. smegmatis* was grown in 7H9 broth containing kanamycin (50 μg/mL) and the fluorescence was measured after 48 h.

To study the activity of *Rv3230c-Rv3229c* promoter in hypoxia, a reporter construct was made using pMV261-*GFP AAV* vector which has GFP-AAV, a short half-life GFP, cloned into *Bam*HI and *Hind*III sites. The target promoter with *Rv1019* ORF was cloned upstream of *GFP-AAV* into *Not*I and *Bam*HI sites by removing the native *hsp60* promoter. Constructs were transformed into *M. smegmatis* strains and the transformants were selected on 7H10 agar containing kanamycin (50 μg/mL). Hypoxia in *M. smegmatis* was induced according to a previously described protocol [35].

### GFP assay

Recombinant *M. smegmatis* containing different promoter constructs was grown until exponential phase, and was then inoculated into fresh medium at an A_600_ of 0.05 and allowed to grow for another 48 h at 37 °C. To analyse the GFP activity during hypoxia, cultures were harvested during respective stages. Absorbance of the culture was measured at 600 nm (A_600_) and fluorescence was measured at the excitation and emission maxima of 488 nm and 520 nm, respectively, in a fluorescent microplate reader (Tecan). Relative Fluorescence Unit (RFU) was determined as measured fluorescence units/A_600_. Background fluorescence due to read-through transcription of the transcriptional fusion vector was determined by measuring the RFU of *M. smegmatis* harbouring empty vector. All measurements were carried out in triplicate. For qualitative analysis, culture was transferred to glass bottomed 96-well plates and observed under a confocal microscope (Nikon A1R-Eclipse Ti, Tokyo, Japan) and the images were analysed with Nikon NIS Elements AR v 4.0.

### Electrophoretic mobility shift assay (EMSA)

EMSA was performed according to Gomez et al [34]. DNA fragments used for EMSA were prepared either by PCR or by annealing complementary oligos. The assays were carried out in 20 µl reactions containing template DNA (5 nM), increasing concentrations of Rv1019 (2-10 µM), 1X buffer containing 10 mM Tris pH 8.0, 1 mM EDTA (pH 8.0), 1mM DTT, 10% glycerol and BSA (100 µg/ml), and were incubated for 30 min at 25 °C. Samples were electrophoresed on 5% or 8% polyacrylamide gel at 100 V at 25 °C, and were stained using ethidium bromide (1µg/ml). For competitive EMSA, 10-fold excess of specific template and for noncompetitive EMSA, 10-fold excess of non-specific template were used.

### Induction of dormancy in *M. tuberculosis* and *M. smegmatis*

Dormancy in *M. tuberculosis* was induced based on Wayne’s model as described by Gopinath *et al* [8]. Briefly, to generate dormant *M. tuberculosis*, we used flat-bottomed glass tubes with a width of 25.5 mm and a liquid holding capacity of 30 ml. Dubos Tween-albumin broth (20 ml) containing 2 × 10^6^ bacteria per milliliter was dispensed into required number of tubes. Cultures were grown with limited internal agitation (130 rpm) using 8-mm Teflon-coated magnetic bars (Sigma-Aldrich, USA) on multipoint magnetic stirrers (Variomag Poly 15, Thermo Scientific, USA). The tubes were closed tightly with caps and wrapped air-tight with parafilm. The whole setup was placed inside a custom-made 37 °C incubator (Santhom Scientific, Bangalore). The status of self-generated hypoxia was monitored visually using methylene blue (final concentration of 1.5 μg/ml). This indicator imparts a greenish blue colour to the culture in the presence of oxygen and turns colourless when oxygen concentration in the medium becomes less than 1%. Growth was monitored every 24 h by measuring A_600_ in a colorimeter (AimilPhotochem, India) and the viability was assessed by plating 100 μL of suitably diluted bacterial culture on 7H10 agar (Difco, USA) in triplicate and incubating at 37 °C for 6-8 weeks.

Dormancy in *M. smegmatis* was induced as described by Dick et al [35]. Briefly, cultures were grown in 20×125 mm test tubes containing 7H9 Middlebrook broth supplemented with Dubos Tween-albumin broth at 37 °C. Seventeen ml of the culture was used throughout and, therefore, the ratio of head space (8.5 ml) to medium (17 ml) was always 1:2. Depending on the conditions of aeration desired, caps with latex liners were either loosely (‘unsealed culture’, continuous oxygen supply) or tightly screwed down (‘sealed culture’, limited oxygen supply). Tubes were stirred using magnetic bars at 150 rpm (utilization of oxygen by the sealed culture leads to its depletion leading to induction of dormancy due to hypoxia). Oxygen depletion was monitored using the oxygen indicator dye methylene blue. Growth and survival of the bacterial populations were monitored by measuring optical density and viable count. Growth was monitored every 24 h by measuring A_600_ in a colorimeter and the viability was assessed by plating 100 μL of suitably diluted bacterial culture on 7H10 agar in triplicate and incubating at 37 °C for 2-3 days.

### RNA isolation, cDNA synthesis and quantitative PCR

RNA was isolated from *Mtb* and *M. smegmatis* using the protocol described by Gopinath et al [8]. Briefly, RNA was extracted using Trizol reagent and precipitated with 100% isopropanol. RNA was resuspended in nuclease-free water and treated with DNaseI. Complementary DNA (cDNA) was prepared using Reverse Transcriptase Core kit (Promega Corporation) according to the manufacturer’s protocol. A qPCR was performed using SYBR Green kit (Takara Bio INC). The thermal cycling protocol was set as follows: an initial denaturation of 95 °C for 1 min; 35 cycles of denaturation at 95 °C for 30 s, primer annealing at 60 °C for 45 s, and extension at 72 °C for 30 s. Following amplification, a melt curve analysis was performed to confirm the specificity of the amplified product. Relative changes in gene expression were calculated using the 2−ΔΔ*C*_t_ method [36], with sigma factor A (*sigA*) as the housekeeping control gene.

### Computational analysis

Mycobaterium database https://mycobrowser.epfl.ch/ was used to collect information on protein and DNA sequences. Protein alignment was performed at NCBI Blastp suite https://blast.ncbi.nlm.nih.gov/Blast.cgi?PAGE=Proteins. Palindromic sequences finder http://www.biophp.org/minitools/find_palindromes/demo.php was used to identify palindromes in the promoter region. MEME was set with a minimum width of 6 bp and a maximum of 50 bp to find palindromic motifs to identify putative binding sites of transcriptional regulators [37] and was set to return a maximum of three motifs. MTB Network Portal database (http://networks.systemsbiology.net/mtb) was used to identify the targets of Rv1019.

## Abbreviations

*Mtb*: *Mycobacterium tuberculosis*
*MSMEG*: *Mycobacterium smegmatis*
TB: tuberculosis
NRP: non-replicating persistence
TetR: tetracycline resistance regulator
ChIP: chromatin immunoprecipitation
EMSA: electrophoretic mobility shift assay.

## Author Contributions

A.R.P and R.A.K designed and conceived the experiments. R.A.K supervised the execution of the experiments. A.R.P and L.K.E performed the experiments. A.R.P and R.A.K wrote the manuscript.

## Acknowledgements

R.A.K thanks Department of Biotechnology, Government of India for financial support. We are grateful to Dr Krishna Kurthkoti for providing plasmids for reporter assays.

## Notes

*Conflicts of interest*: The authors declare that there is no conflict of interest regarding the publication of this article.

### Competing Interest Statement

The authors have declared no competing interest.

## References

1. Westblade, L. F., Campbell, E. A., Pukhrambam, C., Padovan, J. C., Nickels, B. E., Lamour, V. & Darst, S. A. (2010) Structural basis for the bacterial transcription-repair coupling factor/RNA polymerase interaction, Nucleic Acids Res. 38, 8357–69.

2. Organization, W. H. (2021) in Global tuberculosis report 2021 Licence: CC BY-NC-SA 30 IGO

3. Rustad, T. R., Sherrid, A. M., Minch, K. J. & Sherman, D. R. (2009) Hypoxia: a window into Mycobacterium tuberculosis latency, Cell Microbiol. 11, 1151–9.

4. Murphy, D. J. & Brown, J. R. (2007) Identification of gene targets against dormant phase Mycobacterium tuberculosis infections, BMC Infect Dis. 7, 84.

5. Cole, S. T., Brosch, R., Parkhill, J., Garnier, T., Churcher, C., Harris, D., Gordon, S. V., Eiglmeier, K., Gas, S., Barry, C. E., 3rd, Tekaia, F., Badcock, K., Basham, D., Brown, D., Chillingworth, T., Connor, R., Davies, R., Devlin, K., Feltwell, T., Gentles, S., Hamlin, N., Holroyd, S., Hornsby, T., Jagels, K., Krogh, A., McLean, J., Moule, S., Murphy, L., Oliver, K., Osborne, J., Quail, M. A., Rajandream, M. A., Rogers, J., Rutter, S., Seeger, K., Skelton, J., Squares, R., Squares, S., Sulston, J. E., Taylor, K., Whitehead, S. & Barrell, B. G. (1998) Deciphering the biology of Mycobacterium tuberculosis from the complete genome sequence, Nature. 393, 537–44.

6. Rosenkrands, I., Slayden, R. A., Crawford, J., Aagaard, C., Barry, C. E., 3rd & Andersen, P. (2002) Hypoxic response of Mycobacterium tuberculosis studied by metabolic labeling and proteome analysis of cellular and extracellular proteins, J Bacteriol. 184, 3485–91.

7. Eoh, H. & Rhee, K. Y. (2013) Multifunctional essentiality of succinate metabolism in adaptation to hypoxia in Mycobacterium tuberculosis, Proc Natl Acad Sci U S A. 110, 6554–9.

8. Gopinath, V., Raghunandanan, S., Gomez, R. L., Jose, L., Surendran, A., Ramachandran, R., Pushparajan, A. R., Mundayoor, S., Jaleel, A. & Kumar, R. A. (2015) Profiling the Proteome of Mycobacterium tuberculosis during Dormancy and Reactivation, Mol Cell Proteomics. 14, 2160–76.

9. Peterson, E. J., Reiss, D. J., Turkarslan, S., Minch, K. J., Rustad, T., Plaisier, C. L., Longabaugh, W. J., Sherman, D. R. & Baliga, N. S. (2014) A high-resolution network model for global gene regulation in Mycobacterium tuberculosis, Nucleic Acids Res. 42, 11291–303.

10. Dick, T. (2001) Dormant tubercle bacilli: the key to more effective TB chemotherapy?, J Antimicrob Chemother. 47, 117–8.

11. Parrish, N. M., Dick, J. D. & Bishai, W. R. (1998) Mechanisms of latency in Mycobacterium tuberculosis, Trends Microbiol. 6, 107–12.

12. Minch, K. J., Rustad, T. R., Peterson, E. J., Winkler, J., Reiss, D. J., Ma, S., Hickey, M., Brabant, W., Morrison, B., Turkarslan, S., Mawhinney, C., Galagan, J. E., Price, N. D., Baliga, N. S. & Sherman, D. R. (2015) The DNA-binding network of Mycobacterium tuberculosis, Nat Commun. 6, 5829.

13. Pushparajan, A. R., Ramachandran, R., Gopi Reji, J. & Ajay Kumar, R. (2020) Mycobacterium tuberculosis TetR family transcriptional regulator Rv1019 is a negative regulator of the mfd-mazG operon encoding DNA repair proteins, FEBS Lett. 594, 2867–2880.

14. Schubert, O. T., Ludwig, C., Kogadeeva, M., Zimmermann, M., Rosenberger, G., Gengenbacher, M., Gillet, L. C., Collins, B. C., Röst, H. L., Kaufmann, S. H., Sauer, U. & Aebersold, R. (2015) Absolute Proteome Composition and Dynamics during Dormancy and Resuscitation of Mycobacterium tuberculosis, Cell Host Microbe. 18, 96–108.

15. Turkarslan, S., Peterson, E. J., Rustad, T. R., Minch, K. J., Reiss, D. J., Morrison, R., Ma, S., Price, N. D., Sherman, D. R. & Baliga, N. S. (2015) A comprehensive map of genome-wide gene regulation in Mycobacterium tuberculosis, Sci Data. 2, 150010.

16. Betts, J. C., Lukey, P. T., Robb, L. C., McAdam, R. A. & Duncan, K. (2002) Evaluation of a nutrient starvation model of Mycobacterium tuberculosis persistence by gene and protein expression profiling, Mol Microbiol. 43, 717–31.

17. Schnappinger, D., Ehrt, S., Voskuil, M. I., Liu, Y., Mangan, J. A., Monahan, I. M., Dolganov, G., Efron, B., Butcher, P. D., Nathan, C. & Schoolnik, G. K. (2003) Transcriptional Adaptation of Mycobacterium tuberculosis within Macrophages: Insights into the Phagosomal Environment, J Exp Med. 198, 693–704.

18. Wayne, L. G. & Hayes, L. G. (1996) An in vitro model for sequential study of shiftdown of Mycobacterium tuberculosis through two stages of nonreplicating persistence, Infect Immun. 64, 2062–9.

19. Camus, J. C., Pryor, M. J., Médigue, C. & Cole, S. T. (2002) Re-annotation of the genome sequence of Mycobacterium tuberculosis H37Rv, Microbiology (Reading). 148, 2967–2973.

20. Phetsuksiri, B., Jackson, M., Scherman, H., McNeil, M., Besra, G. S., Baulard, A. R., Slayden, R. A., DeBarber, A. E., Barry, C. E., 3rd, Baird, M. S., Crick, D. C. & Brennan, P. J. (2003) Unique mechanism of action of the thiourea drug isoxyl on Mycobacterium tuberculosis, J Biol Chem. 278, 53123–30.

21. Chang, Y., Wesenberg, G. E., Bingman, C. A. & Fox, B. G. (2008) In vivo inactivation of the mycobacterial integral membrane stearoyl coenzyme A desaturase DesA3 by a C-terminus-specific degradation process, J Bacteriol. 190, 6686–96.

22. Ermolaeva, M. D., White, O. & Salzberg, S. L. (2001) Prediction of operons in microbial genomes, Nucleic Acids Res. 29, 1216–21.

23. Deng, W., Li, C. & Xie, J. (2013) The underling mechanism of bacterial TetR/AcrR family transcriptional repressors, Cell Signal. 25, 1608–13.

24. Cuthbertson, L. & Nodwell, J. R. (2013) The TetR family of regulators, Microbiol Mol Biol Rev. 77, 440–75.

25. Park, H. D., Guinn, K. M., Harrell, M. I., Liao, R., Voskuil, M. I., Tompa, M., Schoolnik, G. K. & Sherman, D. R. (2003) Rv3133c/dosR is a transcription factor that mediates the hypoxic response of Mycobacterium tuberculosis, Mol Microbiol. 48, 833–43.

26. Rehberg, N., Omeje, E., Ebada, S. S., van Geelen, L., Liu, Z., Sureechatchayan, P., Kassack, M. U., Ioerger, T. R., Proksch, P. & Kalscheuer, R. (2019) 3-O-Methyl-Alkylgallates Inhibit Fatty Acid Desaturation in Mycobacterium tuberculosis, Antimicrob Agents Chemother. 63.

27. Di Capua, C. B., Doprado, M., Belardinelli, J. M. & Morbidoni, H. R. (2017) Complete auxotrophy for unsaturated fatty acids requires deletion of two sets of genes in Mycobacterium smegmatis, Mol Microbiol. 106, 93–108.

28. Rodríguez, J. G., Hernández, A. C., Helguera-Repetto, C., Aguilar Ayala, D., Guadarrama-Medina, R., Anzóla, J. M., Bustos, J. R., Zambrano, M. M., González, Y. M. J., García, M. J. & Del Portillo, P. (2014) Global adaptation to a lipid environment triggers the dormancy-related phenotype of Mycobacterium tuberculosis, mBio. 5, e01125–14.

29. Sharma, S., Ryndak, M. B., Aggarwal, A. N., Yadav, R., Sethi, S., Masih, S., Laal, S. & Verma, I. (2017) Transcriptome analysis of mycobacteria in sputum samples of pulmonary tuberculosis patients, PLoS One. 12, e0173508.

30. Sikri, K., Batra, S. D., Nandi, M., Kumari, P., Taneja, N. K. & Tyagi, J. S. (2015) The pleiotropic transcriptional response of Mycobacterium tuberculosis to vitamin C is robust and overlaps with the bacterial response to multiple intracellular stresses, Microbiology (Reading). 161, 739–53.

31. Kendall, S. L., Burgess, P., Balhana, R., Withers, M., Ten Bokum, A., Lott, J. S., Gao, C., Uhia-Castro, I. & Stoker, N. G. (2010) Cholesterol utilization in mycobacteria is controlled by two TetR-type transcriptional regulators: kstR and kstR2, Microbiology (Reading). 156, 1362–1371.

32. García-Fernández, E., Medrano, F. J., Galán, B. & García, J. L. (2014) Deciphering the transcriptional regulation of cholesterol catabolic pathway in mycobacteria: identification of the inducer of KstR repressor, J Biol Chem. 289, 17576–88.

33. Kahramanoglou, C., Cortes, T., Matange, N., Hunt, D. M., Visweswariah, S. S., Young, D. B. & Buxton, R. S. (2014) Genomic mapping of cAMP receptor protein (CRP Mt) in Mycobacterium tuberculosis: relation to transcriptional start sites and the role of CRPMt as a transcription factor, Nucleic Acids Res. 42, 8320–9.

34. Gomez, R. L., Jose, L., Ramachandran, R., Raghunandanan, S., Muralikrishnan, B., Johnson, J. B., Sivakumar, K. C., Mundayoor, S. & Kumar, R. A. (2016) The multiple stress responsive transcriptional regulator Rv3334 of Mycobacterium tuberculosis is an autorepressor and a positive regulator of kstR, Febs j. 283, 3056–71.

35. Dick, T., Lee, B. H. & Murugasu-Oei, B. (1998) Oxygen depletion induced dormancy in Mycobacterium smegmatis, FEMS Microbiol Lett. 163, 159–64.

36. Livak, K. J. & Schmittgen, T. D. (2001) Analysis of relative gene expression data using real-time quantitative PCR and the 2(−Delta Delta C(T)) Method, Methods. 25, 402–8.

37. Bailey, T. L., Boden, M., Buske, F. A., Frith, M., Grant, C. E., Clementi, L., Ren, J., Li, W. W. & Noble, W. S. (2009) MEME SUITE: tools for motif discovery and searching, Nucleic Acids Res. 37, W202–8.

