## Supplementary figures and tables for "*Mycobacterium tuberculosis* transcriptional regulator Rv1019 is upregulated in hypoxia and negatively regulates *Rv3230c*-*Rv3229c* operon encoding enzymes in the oleic acid biosynthetic pathway"

### Supporting Information

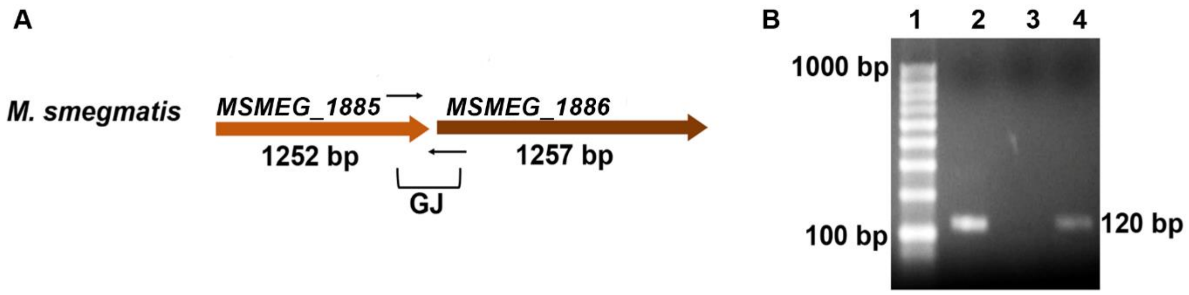

**Fig. S1 Cotranscription of *MSMEG_1885* (≈*Rv3230c*) and *MSMEG_1886* (≈*Rv3229c*) in *M. smegmatis*. (A)** Representative orientation and size of *MSMEG_1885* and *MSMEG_1886* in *M. smegmatis*. Gene junctions are indicated (GJ). Thin black arrows represent the DNA region selected to design PCR primers to amplify the gene junctions; (**B**) RT-PCR amplification to prove cotranscription of *MSMEG_1885* and *MSMEG_1886*. Lane 1: 100 bp DNA ladder, Lane 2: positive control, Lane 3: PCR amplification from negative reverse transcription reaction, Lane 4: PCR amplification from positive reverse transcription reaction.

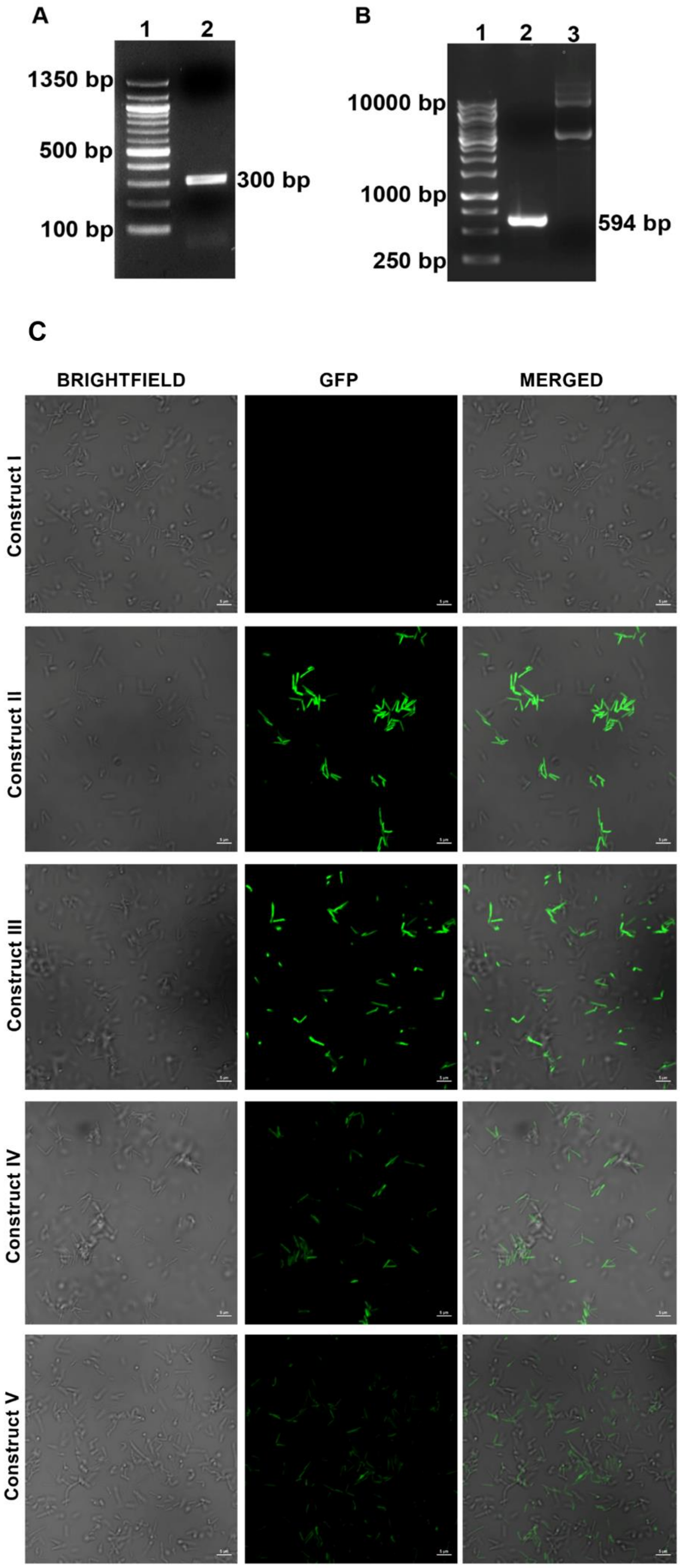
A
1
2
1350 bp
500 bp
100 bp
300 bp
B
1
2
3
10000 bp
1000 bp
250 bp
594 bp
C
BRIGHTFIELD
GFP
MERGED
Construct I
Construct II
Construct III
Construct IV
Construct V

**Fig. S2 Reporter assay in *M. smegmatis*. (A)** PCR amplification of upstream 300-bp region of *Rv3230c* ORF. Lane 1: 100 bp DNA ladder, Lane 2: PCR amplicon of *Rv3230c* promoter; (**B**) Cloning of *Rv1019* ORF inpFPV27 vector. Lane 1: 1 kb DNA ladder, Lane 2: PCR amplicon of *Rv1019* ORF, Lane 3: pFPV27 vector; (**C**) Confocal photomicrographs of *M. smegmatis* Δ*MSMEG_5424* carrying constructs I, II, III, IV or V. Scale bars are 5 µm.

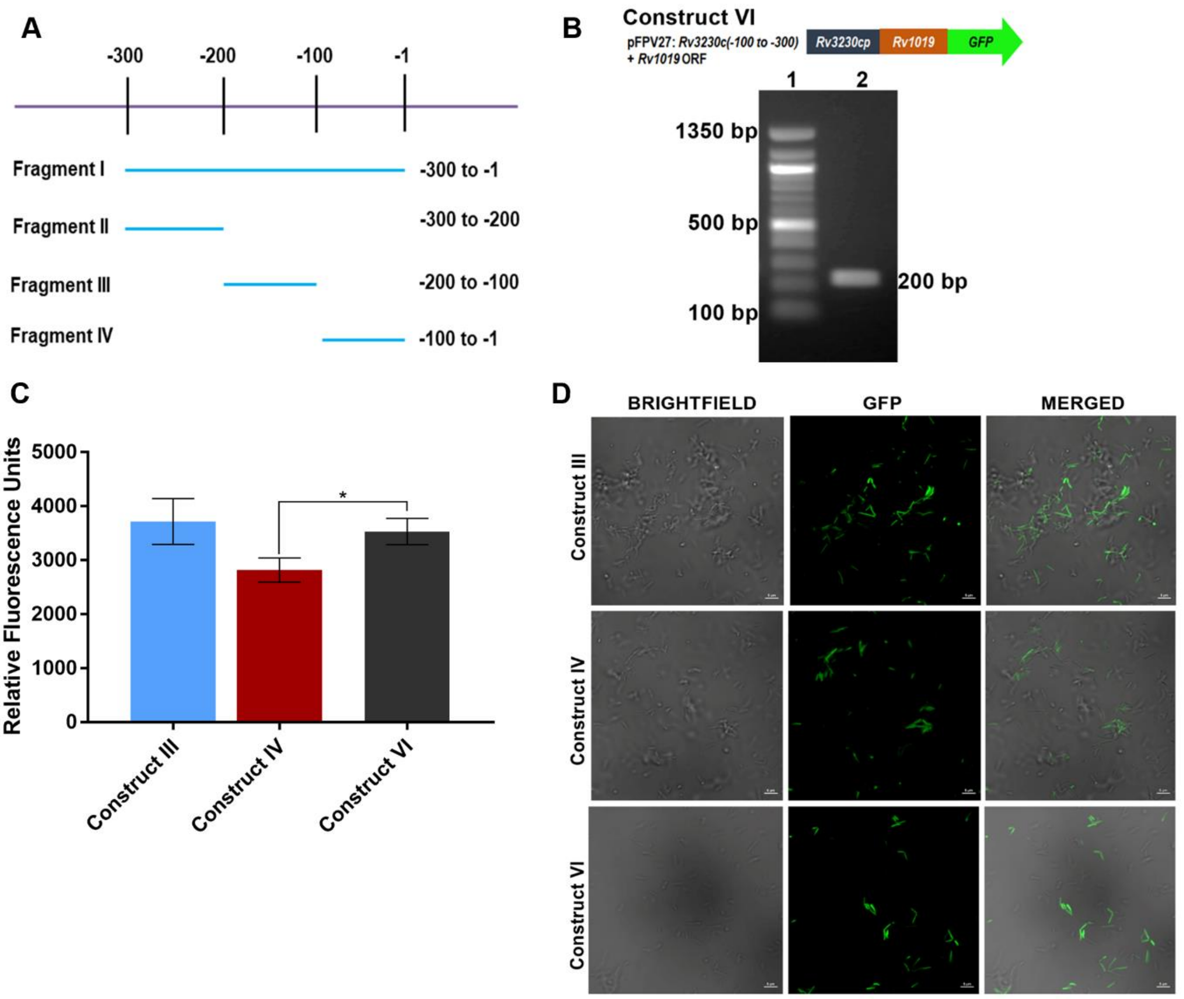

**Fig. S3 Validation of binding sites of Rv1019 on *Rv3230c-Rv3229c* promoter. (A)** Representative image showing fragments used to map the binding region upstream of *Rv3230c* ORF; **(B)**PCR amplification of *Rv1019* ORF and *Rv3230c-Rv3229c* promoter (without 12-bp binding region). Lane 1: 100 bp DNA ladder, Lane 2: *Rv3230c-Rv3229c* promoter (-100 to -300 bp region without 12-bp cognate binding sequence). Construct VI: -100 to -300 region with *Rv1019* ORF was cloned upstream of GFP in pFPV27 vector. **(C)** GFP expression from construct IV vs Construct VI was compared and represented in terms of RFU. The values are the mean ± standard deviation of three independent experiments (Tukey's multiple comparisons test). *P ≤ 0.02. **(D)** Confocal photomicrographs of *M. smegmatis* carrying constructs III, IV and VI. Scale bars are 5 μm.

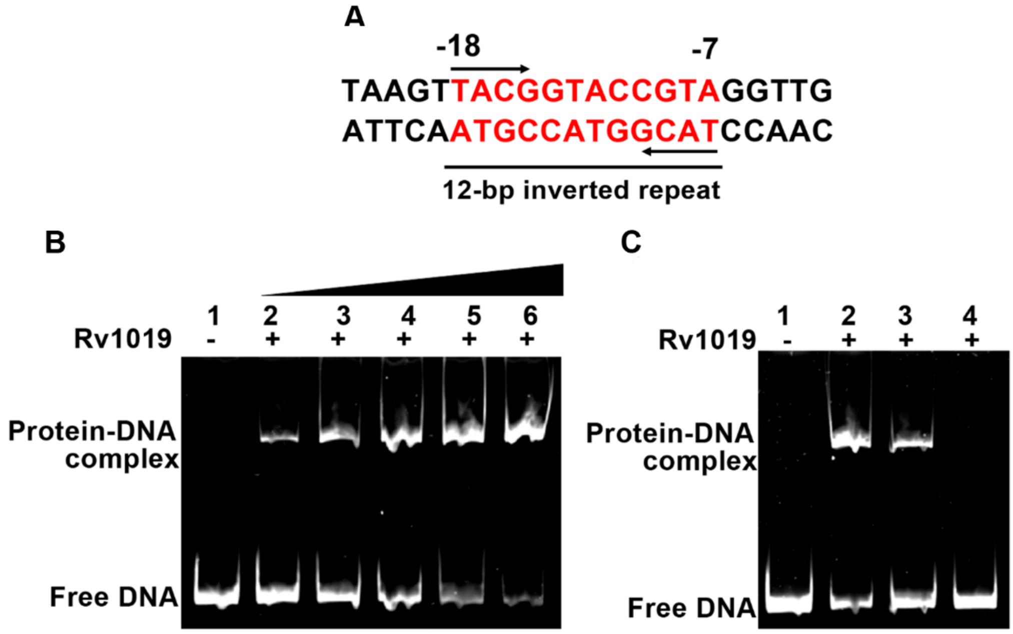

**Fig. S4 Rv1019 binding site is conserved in *M. smegmatis***. (**A**) Representative image of 12-bp inverted repeat present on *M. smegmatis MSMEG_1885(≈Rv3230c*). Inverted repeat (red) and flanking region (black); (**B**) EMSA on 12-bp palindrome of *M. smegmatis*. Lane 1: palindromic DNA sequence without Rv1019, Lanes 2–6: DNA with increasing concentrations (2-10 μM) of Rv1019; **(C)** Competitive and noncompetitive EMSA to validate the specificity of binding of Rv1019 on *M. smegmatis* palindromic sequence. Lane 1: DNA without protein, Lane 2: with DNA incubated with 10 μM of Rv1019 (positive control), Lane 3: competitive EMSA, Lane 4: noncompetitive EMSA.

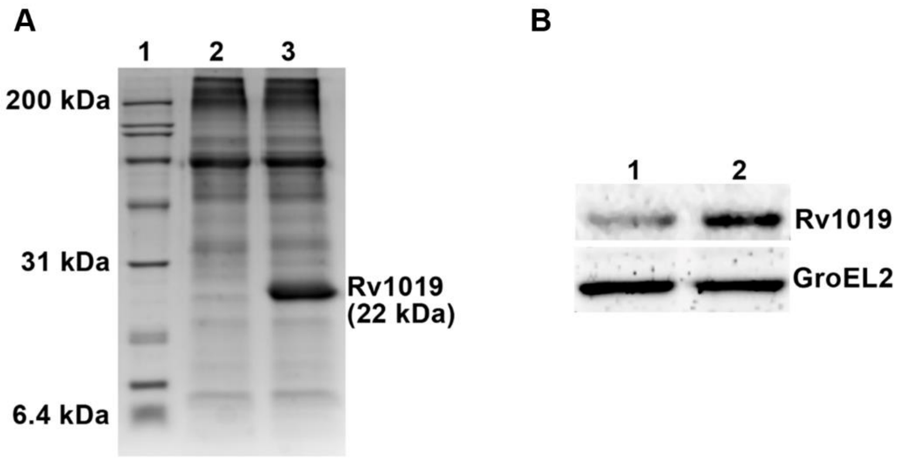

**Fig. S5 Constitutive expression of *Rv1019* in *M. tuberculosis*. (A)** SDS-PAGE shows constitutive expression of *Rv1019* in *M. tuberculosis*. Lane 1: broad range protein ladder, Lane 2: vector control (*M. tuberculosis:pBEN*), Lane 3: *M. tuberculosis:pBEN:Rv1019*. (**B**) Western blotting shows the constitutive expression of *Rv1019*. Lane 1: vector control (*M. tuberculosis: pBEN*), Lane 2: *M. tuberculosis: pBEN:Rv1019*.

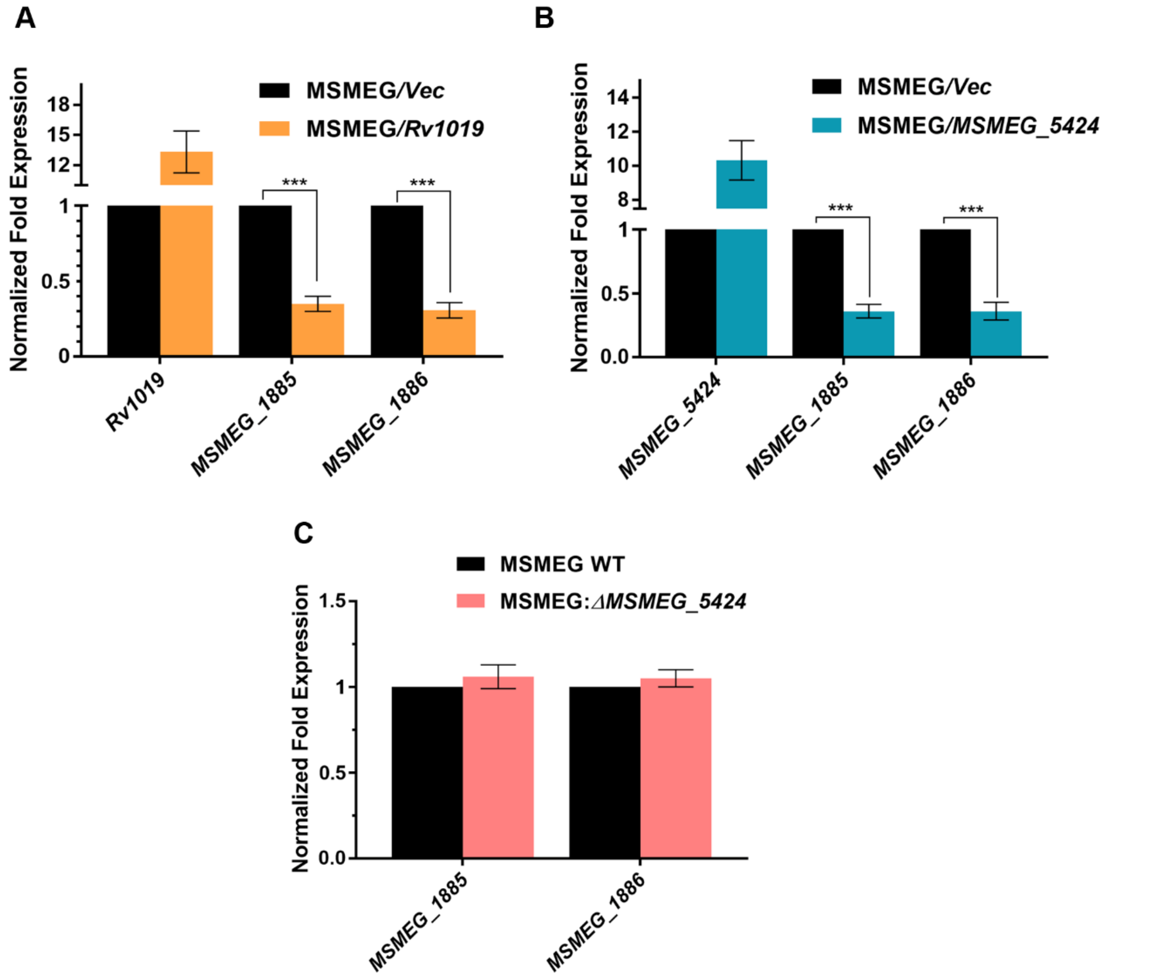

**Fig. S6 Analysis of expression of *MSMEG_1885* and *MSMEG_1886* in *M. smegmatis*.** (**A**) Quantitative real-time PCR showing expression of *Rv1019* (constitutive), and the expression of endogenous *MSMEG_1885*and *MSMEG_1886* in *M. smegmatis*; (**B**) Quantitative real-time PCR showing expression of *MSMEG_5424* (constitutive), and the expression of endogenous *MSMEG_1885* and *MSMEG_1886*in *M. smegmatis*. ***P ≤ 0.0001; (**C**) Quantitative real-time PCR showing expression of *MSMEG_1885*and *MSMEG_1886* in *ΔMSMEG_5424* mutant of *M. smegmatis*. Values are the mean ± standard deviation of three independent experiments (Student's *t*-test).

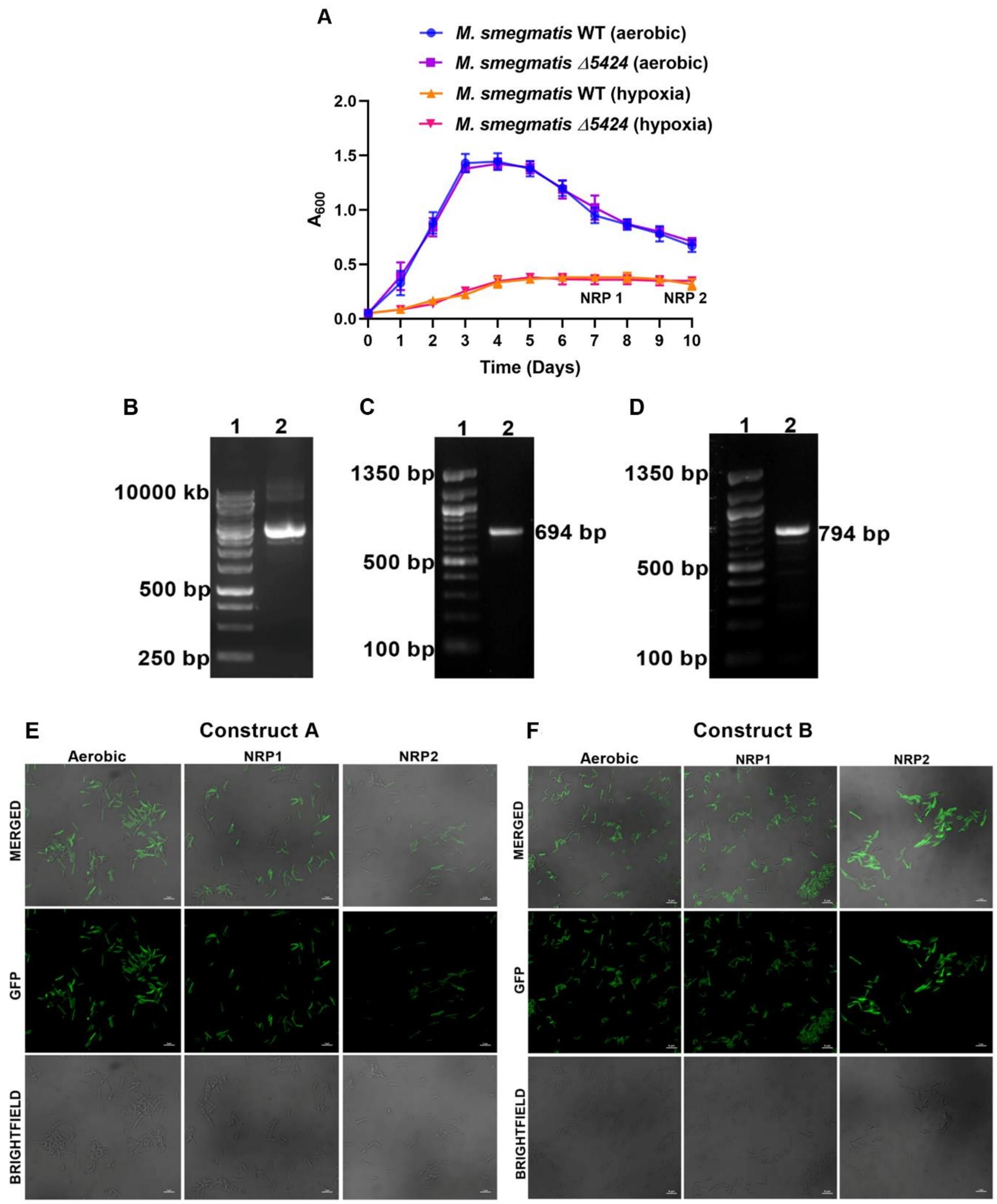

**Fig. S7 GFP Reporter assay in *M. smegmatis* under hypoxia.** (**A**) Measurement of growth of *M. smegmatis* by optical density. (Values at each point of time represent an average value from three tubes); (**B**) Agarose gel analysis of pMV261:*GFP-AAV* vector. Lane 1: 1 kb DNA ladder, Lane 2: pMV261:*GFP-AAV* vector; (**C**) Construct A: cloning of 12-bp cognate binding site of Rv1019 on *Rv3230c-Rv3229c* promoter (including 5-bp upstream and downstream flanking regions) and *Rv1019* ORF (594 bp) into pMV261:*GFP-AAV*. Lane 1: 100 bp DNA ladder, Lane 2: PCR amplification of 12-bp cognate binding site with flanking regions and *Rv1019* ORF from Construct VI (Fig. S3 B); (**D**) Construct B: cloning of 18-bp cognate binding site of Rv1019 on its own promoter (including 5-bp upstream and downstream flanking regions) and *Rv1019* ORF into pMV261:*GFP-AAV*. Lane 1: 100 bp DNA ladder, Lane 2: PCR amplification of *Rv1019* ORF with 18-bp cognate binding site on its own promoter; (**E**) Confocal photomicrography of *M. smegmatis* Δ*MSMEG_5424* harbouring Construct A. GFP expression from *M. smegmatis* Δ*MSMEG_5424* harbouring Construct A during aerobic growth, NRP1 and NRP2; (**F**) Confocal photomicrography of *M. smegmatis* Δ*MSMEG_5424* harbouring Construct B. GFP expression from *M. smegmatis* Δ*MSMEG_5424* harbouring Construct B during aerobic growth, NRP1 and NRP2. Scale bars are 5 µm.

**Table 1**

**List of bacterial strains and plasmids used in this study**

| **Strain or plasmid or primers** | **Description** | **Reference or source** |
|---|---|---|
| **Strains** | | |
| *E. coli* JM1019 | *traD36 proA*$^+$*B*$^+$ *lacIq Δ(lacZ)M15/ Δ(lac-proAB) glnV44 e14- gyrA96 recA1 relA1 endA1 thi hsdR17* | Invitrogen, California, USA |
| *Mycobacterium tuberculosis* H37Rv | Wild type | NIRT, Chennai, India |
| *Mycobacterium smegmatis* mc$^2$155 | Wild type | NIRT, Chennai, India |
| *Mycobacterium smegmatis* mc$^2$155:*ΔMSMEG_5424* | *MSMEG_5424* deletion mutant of *Mycobacterium smegmatis* mc$^2$155 | Pushparajan *et al*., 2020 |
| *Mycobacterium smegmatis* mc$^2$155:*Rv1019* | *Rv1019* overexpressing *Mycobacterium smegmatis* mc$^2$155 | Pushparajan *et al*., 2020 |
| *Mycobacterium smegmatis* mc$^2$155:*MSMEG_5424* | *MSMEG_5424*overexpressing *Mycobacterium smegmatis* mc$^2$155 | Pushparajan *et al*., 2020 |

| | | |
|---|---|---|
| **Plasmids** | | |
| pFPV27 | Kan$^r$; promoter-less reporter system | Lalitha Ramakrishnan, currently at University of Cambridge, UK |
| pFPV60 | Kan$^r$; *Hsp60* promoter reporter system | Lalitha Ramakrishnan |
| pBEN | Kan$^r$; *Hsp60* promoter mycobacterial shuttle vector | Lalitha Ramakrishnan |
| pMV261:*GFP-AAV* | Kan$^r$; *Hsp60* promoter mycobacterial shuttle vector containing *GFP-AAV* | Krishna Kurthkoti |

**Table 2**

**List of primers and oligos used in this study**

| **Primer / Oligo Name** | **Sequence (5’ --------------- 3’)** | **Description** |
|---|---|---|
| pBEN-*Rv1019* F | CGCggatccATGACGGGGACCGAGCGC | *Rv1019* amplification primer to clone into pBEN, *Bam*HI site (lower case). Forward primer. |
| pBEN-*Rv1019* R | CCCAAGCTTCTACTCGTCCTGTAGCCGCGG | *Rv1019* amplification primer to clone into pBEN, *Hind*III site (lower case). Reverse primer. |
| *Hsp60* Pro F | CGGTCATGGGCCGAACATAC | *Hsp60* promoter 300 bp upstream amplification primer. Forward primer. |
| *Hsp60* Pro R | TGCGAAGTGATTCCTCCGGATCGG | *Hsp60* promoter 300 bp upstream amplification primer. Reverse primer. |
| T4g32 S1 | GGGACCCTAGAGGTCCCCTTT | Transcription terminator sequence, strand 1 |
| T4g32 S2 | AAAGGGGACCTCTAGGGTCCC | Transcription terminator sequence, strand 2 |
| *Sig A*RTF | AGGATCAACTGCAGTCGGTG | qRT-PCR primer for sigma factor A. Forward primer. |
| *Sig A*RTR | CTTGGATTCGATCTGGCGGA | qRT-PCR primer for sigma factor A. Reverse primer. |
| *Rv1019* RT F | ATTCTGGCTGGAGACTTCGC | qRT-PCR primer for *Rv1019*. Forward primer. |

| | | |
|---|---|---|
| *Rv1019* RT R | GCCATTCCAGACCAGGTTGA | qRT-PCR primer for *Rv1019*. Reverse primer. |
| *MSMEG_5424* (*Rv1019)* RT F | GCGACGCTGCTCAATGAC | qRT-PCR primer for *MSMEG_5424*. Forward primer. |
| *MSMEG_5424* (*Rv1019)* RT R | CATGGACACCGAACCGAC | qRT-PCR primer for *MSMEG_5424*. Reverse primer. |
| pFPV-*Rv3230c* Pro F | CGCggatccGGGTGTGACGCAGTCTGCGC | *Rv3230c* 300 bp upstream promoter amplification primer to clone into pFPV27, *Bam*HI site (lower case). Forward primer. |
| pFPV-*Rv3230c* Pro R | CCGgaattcAGGAAGCTCCTGCTCGGCCT | *Rv3230c* 300 bp upstream promoter amplification primer to clone into pFPV27, *Eco*RI site (lower case). Reverse primer. |
| pFPV-*Rv1019* ORF F | CCGgaattcATGACGGGGACCGAGCGCC | *Rv1019* ORF amplification primer to clone into pFPV27, *Eco*RI site (lower case). Forward primer. |
| pFPV-*Rv1019* ORF R | GATCggtaccCTACTCGTCCTGTAGCCGCGG | *Rv1019* ORF amplification primer to clone into pFPV27, *Kpn*I site (lower case). Reverse primer. |
| *Rv3230c* -*Rv3229c* Pro FR I F | GGGTGTGACGCAGTCTGCGC | EMSA fragment I primer, *Rv3230c* -*Rv3229c*upstream -300 to -1 bp. Forward primer. |
| *Rv3230c* -*Rv3229c* Pro FR I R | AGGAAGCTCCTGCTCGGCCT | EMSA fragment I primer, *Rv3230c* -*Rv3229c*upstream -300 to -1 bp. Reverse primer. |
| *Rv3230c* -*Rv3229c* Pro FR II F | GGGTGTGACGCAGTCTGCGC | EMSA fragment II primer, *Rv3230c* -*Rv3229c*upstream -300 to -200 bp. Forward primer. |
| *Rv3230c* -*Rv3229c* Pro FR II R | TCGTCGCCCAGGTCAGCGGC | EMSA fragment II primer, *Rv3230c* -*Rv3229c*upstream -300 to -200 bp. Reverse primer. |
| *Rv3230c* -*Rv3229c* Pro FR III F | ACGGCATCGTCAGAGCAGGT | EMSA fragment III primer, *Rv3230c* -*Rv3229c*upstream -200 to -100 bp. Forward primer. |
| *Rv3230c* -*Rv3229c* Pro FR III R | ACGGCATCGTCAGAGCAGGT | EMSA fragment III primer, *Rv3230c* -*Rv3229c*upstream -200 to -100 bp. Reverse primer. |
| *Rv3230c* -*Rv3229c* Pro FR IV F | GGGTGTGACGCAGTCTGCGC | EMSA fragment IV primer, *Rv3230c* -*Rv3229c*upstream -100 to -1 bp. Forward primer. |
| *Rv3230c* -*Rv3229c* Pro FR IV R | AGGAAGCTCCTGCTCGGCCT | EMSA fragment IV primer, *Rv3230c* -*Rv3229c*upstream -100 to -1 bp. Reverse primer. |
| pFPV-*Rv3230c* -*Rv3229c* Upstream w/o ProF | CGCggatccCGATCGATCAGGTTGTCGCC | *Rv3230c* -*Rv3229c* upstream primer for amplifying region without binding site to clone into pFPV27, *Bam*HI site (lower case). Forward primer. |

| | | |
|---|---|---|
| pFPV-*Rv3230c* -*Rv3229c* Upstream w/o ProR | CCGgaattcACGGCATCGTCAGAGCAGGT | *Rv3230c* -*Rv3229c* upstream primer for amplifying region without binding site to clone into pFPV27, *Eco*RI site (lower case). Reverse primer. |
| *Rv3230c* -*Rv3229c* GP F | ATTGCGTGCTGGACATCTAG | Junction primer to amplify *Rv3230c* -*Rv3229c*. Forward primer. |
| *Rv3230c* -*Rv3229c* GP R | ACGTCGACGTCAGTGATCGC | Junction primer to amplify *Rv3230c* -*Rv3229c*. Reverse primer. |
| *MSMEG_1885-MSMEG_1886* GP F | GACTGTGTGCTCGATGTTTAG | Junction primer to amplify *MSMEG_1885-MSMEG_1886*. Forward primer. |
| *MSMEG_1885-MSMEG_1886* GP R | GCGCGAACGCCGGGACGTCG | Junction primer to amplify *MSMEG_1885-MSMEG_1886*. Reverse primer. |
| *Rv3230c* RT F | GCTTAACGCCAGCATCATC | qRT-PCR primer for *Rv3230c*. Forward primer. |
| *Rv3230c* RT R | GTAGTCGAAACTGAAGCCC | qRT-PCR primer for *Rv3230c*. Reverse primer. |
| *Rv3229c* RT F | ACCACAAATACACCAACATCC | qRT-PCR primer for *Rv3229c*. Forward primer. |
| *Rv3229c* RT R | CGACATAGTCCTTGAACACC | qRT-PCR primer for *Rv3229c*. Reverse primer. |
| *Rv3230c* PAL S1 | taggtTACGGTACCGTAgcttt | Rv1019 cognitive binding sequence on *Rv3230c-Rv3229c* promoter (in uppercase), strand 1 |
| *Rv3230c* PAL S2 | tttcgATGCCATGGCATtggat | Rv1019 cognitive binding sequence on *Rv3230c-Rv3229c* promoter (in uppercase), strand 2 |
| *MS_1885-MS_1886* PAL S1 | taagtTACGGTACCGTAggttg | cognitive binding sequence on *MSMEG_1885-MSMEG_1886* promoter (in uppercase), strand 1 |
| *MS_1885-MS_1886* PAL S2 | attcaATGCCATGGCATccaac | cognitive binding sequence on *MSMEG_1885-MSMEG_1886* promoter (in uppercase), strand 2 |
| Random oligo S1 | GATCGATCGATCGATCGA | Non-specific sequence from *Hsp60* promoter, strand 1 |
| Random oligo S2 | CTAGCTAGCTAGCTAGCT | Non-specific sequence from *Hsp60* promoter, strand 2 |
| *MS_1885* RT F | AACTTCGTGATGCCCGATCC | qRT-PCR primer for *MSMEG_1885*. Forward primer. |
| *MS_1885* RT R | TCAGCTCCGATCCGAACAAC | qRT-PCR primer for *MSMEG_1885*. Reverse primer. |

| | | |
|---|---|---|
| *MS_1886* RT F | CACCCACAACTTCATGCAC | qRT-PCR primer for *MSMEG_1886*. Forward primer. |
| *MS_1886* RT R | AATCCTTGAACACCTGCCC | qRT-PCR primer for *MSMEG_1886*. Reverse primer. |
| pMV-*Rv3230c* -*Rv3229c* Pro + *Rv1019* ORF F | ATAAGAATgcggccgcGGGTGTGACGCAGTCTGCGC | *Rv3230c* -*Rv3229c* promoter and *Rv1019* ORF amplifying primer to clone into pMV261-*GFPAAV*, *Not*I site (lower case). Forward primer. |
| pMV-*Rv3230c* -*Rv3229c* Pro + *Rv1019* ORF R | CGCggatccCTACTCGTCCTGTAGCCGCGG | *Rv3230c* -*Rv3229c* promoter and *Rv1019* ORF amplifying primer to clone into pMV261- *GFPAAV*, *Bam*HI site (lower case). Reverse primer. |
| pMV-*Rv1019* Pro + *Rv1019* ORF F | ATAAGAATgcggccgcTTGGTTCAGATAGTTCAGGTTCG | *Rv1019* promoter + *Rv1019* ORF amplifying primer to clone into pMV261-*GFPAAV*, *Not*I site (lower case). Forward primer. |
| pMV-*Rv1019* Pro + *Rv1019* ORF R | CGCggatccCTACTCGTCCTGTAGCCGCGG | *Rv1019* promoter + *Rv1019* ORF amplifying primer to clone into pMV261-*GFPAAV*, *Not*I site (lower case). Reverse primer. |
